# The role of glutathione in periplasmic redox homeostasis and oxidative protein folding in *Escherichia coli*

**DOI:** 10.1101/2023.03.07.531512

**Authors:** Lisa R. Knoke, Jannik Zimmermann, Natalie Lupilov, Jannis F. Schneider, Beyzanur Celebi, Bruce Morgan, Lars I. Leichert

## Abstract

The thiol redox balance in the periplasm of *E. coli* depends on the DsbA/B pair for oxidative power and the DsbC/D system as its complement for isomerization of non-native disulfides. While the standard redox potentials of those systems are known, the *in vivo* redox potential imposed onto protein thiol disulfide pairs in the periplasm remains unknown. Here, we used genetically encoded redox probes (roGFP2 and roGFP-iL), targeted to the periplasm, to directly probe the thiol redox homeostasis in this compartment. These probes contain two cysteine residues, that are virtually completely reduced in the cytoplasm, but once exported into the periplasm, can form a disulfide bond, a process that can be monitored by fluorescence spectroscopy. Even in the absence of DsbA, roGFP2, exported to the periplasm, was fully oxidized, suggesting the presence of an alternative system for the introduction of disulfide bonds into exported proteins. However, the absence of DsbA shifted the periplasmic thiol-redox potential from -228 mV to a more reducing -243 mV and the capacity to re-oxidize periplasmic roGFP2 after a reductive pulse was significantly decreased. Re-oxidation in a DsbA strain could be fully restored by exogenous oxidized glutathione (GSSG), while reduced GSH accelerated re-oxidation of roGFP2 in the WT. In line, a strain devoid of endogenous glutathione showed a more reducing periplasm, and was significantly worse in oxidatively folding PhoA, a native periplasmic protein and substrate of the oxidative folding machinery. PhoA oxidative folding could be enhanced by the addition of exogenous GSSG in the WT and fully restored in a Δ*dsbA* mutant. Taken together this suggests the presence of an auxiliary, glutathione-dependent thiol-oxidation system in the bacterial periplasm.

## Introduction

Extracellular proteins are stabilized by structural disulfide bonds that are introduced during oxidative protein folding. Thus, the correct folding of most of the proteins translocated into the periplasm and beyond in the model bacterium *Escherichia coli* depends on the formation of native disulfide bonds. The major pathway for introducing disulfide bonds in the periplasm of *E. coli* involves the DsbA-DsbB thiol oxidase system (Bardwell et al., 1993, 1991; Collet and Bardwell, 2002; Missiakas et al., 1993; Wunderlich and Glockshuber, 1993). DsbA contains a thioredoxin-like domain including an active site CxxC motif with a standard redox potential of around –122 mV (Martin et al., 1993; Wunderlich et al., 1993). After disulfide introduction, DsbA itself is re-oxidized by the integral membrane protein DsbB, which in turn is connected to the ubiquinone pool of the bacterial plasma membrane (Christensen et al., 2016; Inaba et al., 2009; Inaba and Ito, 2008; Manta et al., 2019; Vivian et al., 2009). As some proteins contain multiple cysteine residues, cells require quality control systems to isomerize non-native disulfide bonds. In *E. coli*, this system includes the disulfide isomerase DsbC/G and the reductive power from DsbD, which itself is coupled to the cytoplasmic NADPH pool through Thioredoxin A (TrxA) (Andersen et al., 1997; Cho and Collet, 2013; Missiakas et al., 1994; Rietsch et al., 1997; Shevchik et al., 1994; Sone et al., 1997).

In eukaryotes, oxidative folding in the endoplasmic reticulum (ER) is catalyzed in a similar manner by protein disulfide isomerase(s) (PDI), which also contains thioredoxin-like domains. Unlike DsbA, PDI not only acts as thiol oxidase but also functions as a disulfide reductase/isomerase (Ali Khan and Mutus, 2014; Ushioda and Nagata, 2019). PDI is re-oxidized for example by the sulfhydryl oxidase Endoplasmic reticulum oxidoreductin 1 (ERO1-Lα) (Tu and Weissman, 2004).

The tripeptide glutathione (GSH) is the most abundant low molecular weight thiol in many domains of life ranging from bacteria to eukaryotes (Fahey et al., 1978; Smirnova and Oktyabrsky, 2005; Zechmann et al., 2011) and usually it is noted for its reductive properties, keeping the cytoplasm reduced. In its reducing role, glutathione is oxidized to glutathione disulfide (GSSG) (Aslund et al., 1994; Carmel-Harel and Storz, 2000; Masip et al., 2006). In *E. coli*, cytosolic GSH levels have been described to be in the millimolar range (up to 10 mM). The GSH:GSSG ratio ranges from 50:1 to 200:1 and is maintained by the glutathione reductase Gor, dependent on the cellular NADPH-pool (Åslund et al., 1999; Fahey et al., 1978; Greer and Perham, 1986; Masip et al., 2006; Meister, 1988; Tuggle and Fuchs, 1985).

Exponentially growing *E. coli* cells extensively secrete GSH into the culture medium. Several ABC-transporters mediate GSH transport across the inner cell membrane and outer membrane porins, allow GSH or GSSG to enter or leave the periplasm with around 10 % of all synthesized GSH being secreted. (Holyoake et al., 2016; Pittman et al., 2005; Smirnova et al., 2012; Suzuki et al., 2005; Wang et al., 2018). Although glutathione synthesis is restricted to the cytosol, it is exported into the periplasm of *E. coli* and periplasmic glutathione probably accounts for 10–30% of total cellular GSH (Eser et al., 2009; Pittman et al., 2005; Smirnova et al., 2012). It has been suggested that the GSH:GSSG ratio in the periplasm reflects the GSH:GSSG ratio of the culture medium, being approximately 16:1 (Eser et al., 2009; Pittman et al., 2005; Smirnova et al., 2012, 2020, 2005; Song et al., 2021).

Since redox processes in the cell are highly regulated, the presence of glutathione in the periplasm at such high levels strongly indicates that this molecule also plays a role in the redox homeostasis of the periplasm, similar to its role in the ER (Birk et al., 2013; Delaunay-Moisan et al., 2017). In fact, impaired GSH synthesis or reduced GSH import into the periplasm is connected to several phenotypes, similar to those found in cells lacking the DsbA/DsbB- or DsbC-system (Fabianek et al., 2000; Pittman et al., 2005). However, a direct involvement in disulfide bond formation and redox homeostasis for GSH in the periplasm has not yet been established.

Here, we explore the role of GSH in oxidative protein folding and redox homeostasis in the periplasm of *E. coli* by targeting genetically encoded redox-sensitive probes with an engineered disulfide bond to this compartment and analyzing the periplasmic redox dynamics in the presence or absence of DsbA and/or GSH. We provide evidence that GSH indeed plays a key role in disulfide bond formation and redox homeostasis in the periplasm: We observed that GSSG can complement for the loss of DsbA and, unexpectedly, that GSH accelerates disulfide bond formation in the presence of DsbA. In line, the periplasmic redox balance of GSH-deficient cells was slightly shifted to a more reducing redox potential and the oxidative folding of native DsbA substrates in these cells is impaired.

## Materials and methods

### Strains, Plasmids and growth conditions

Bacterial strains and plasmids used in this study are listed in Supplementary tables S1 and S2. *Escherichia coli* DH5α served as host for plasmid construction and storage and *E. coli* BL21 (DE3) was used for DsbA and *E. coli* MG1655 for roGFP2 recombinant protein production. All *E. coli* strains used in this study were routinely cultivated at 37 °C in Luria-Bertani (LB) medium, supplemented with antibiotics when required for plasmid maintenance and marker selection (ampicillin 200 µg/mL or kanamycin 100 µg/mL), if not stated differently. Protein expression in BL21 was induced with 0.4 mM IPTG when the cultures reached an OD_600_ of ∼0.6–0.8 at 37 °C in LB medium and then incubated for 20 h at 20 °C after induction.

*E. coli* mutants from the KEIO collection were used for construction of double deletion strains and analysis of the periplasmic redox state (Baba et al., 2006). For expression of the redox probes, mutant strains (Suppl. Table 1) harboring roGFP-based sensor protein-coding plasmids (pPT_*roGFP2* or pPT_*roGFP-iL*, Supplementary table S2) were cultivated in MOPS minimal medium (Technova, Hollister, CA, USA) at 37 °C until an optical density (OD) of ∼0.5–0.8 was reached. Sensor protein expression was induced with 0.2 mM IPTG and cells were cultivated for 16 h at 20 °C.

### Construction of different roGFP and DsbA expression vectors

For construction of the pPT plasmid enabling periplasmic targeting, the *torA* sequence was PCR amplified using appropriate primers (Supplementary table S3) from genomic *E. coli* DNA (strain MC4100) and cloned into the pCC plasmid (Nilewski et al., 2021) using the introduced *Nde*I 3’ and 5’ restriction sites. The *Nde*I restriction site upstream of *torA* was then removed by QuickChange mutagenesis (Agilent, Waldbronn, Germany) according to the manufacturer’s protocol using primers QC-*Nde*I-fw and QC-*Nde*I-rv (Supplementary table S3) resulting in the pPT plasmid.

In order to target roGFP2 into the periplasm, *roGFP2* was PCR amplified using the primer pair listed in supplementary table S3 from the pCC_*roGFP2* plasmid as template. The introduced *Nde*I and *Eco*RI restriction sites were used to clone the gene into the pPT-plasmid resulting in pPT_*roGFP2* coding for a TorA_roGFP2 fusion protein.

The *roGFP-iL* gene was purchased from Addgene (Watertown, MA, USA) in a pQE30 plasmid harboring an ampicillin resistance gene. The *roGFP-iL* gene was PCR amplified (Primers are listed in Supplementary table S3) with simultaneous introduction of *Xho*I and *EcoR*I restriction sites. The amplified gene was cloned into the pPT plasmid using the introduced restriction sites resulting in the plasmid pPT_*roGFP-iL* (pLK9).

For expression of a roGFP2-*Ec*DsbA(ΔSP) fusion protein, a truncated variant of the *dsbA* (Δ*nt1-48*) gene was PCR amplified lacking the periplasmic sequence (ΔSP). The PCR fragment was cut with *EcoR*I and *Hind*III. AtPrxA-ΔCP was removed from the p416TEF_roGFP2-AtPrxA-ΔCP (Zimmermann et al., 2021) plasmid by the same restriction enzymes and *Ec*DsbA(ΔSP) was cloned into the plasmid backbone, resulting in the plasmid pLK16.

### Construction of E. coli dsbA, gshA double deletion strain by P1 transduction

For the construction of an *E. coli gshA*, *dsbA* double mutant, the kanamycin cassette was removed from the *gshA* KEIO mutant using the plasmid 709-FLPE as indicated by the supplier (Gene Bridges, Heidelberg, Germany). Removal of the kanamycin cassette was verified by colony PCR using *k1, k2* and *gshA* primers listed in supplementary table S3. The *dsbA* mutant from the KEIO collection (strain: JW3832) was used as P1 donor. P1 transduction was carried out as previously described (Thomason et al., 2007). Correct insertion of *dsbA* deletion in marker free *ΔgshA* was checked using appropriate primers (Supplementary table S3).

Periplasmic roGFP2-based measurements in E. coli

For determining the roGFP2 redox state in different *E. coli* WT, Δ*dsbA*, Δ*gshA* or Δ*dsbA*Δ*gshA*, the cells were transformed with the pPT_roGFP2 plasmid encoding the TorA(SP)-roGFP2 fusion construct. Correct periplasmic probe localization was verified by fluorescence microscopy (Supplementary figure 1). After expression of roGFP2 for 16 h at 20 °C, as aforementioned, cells were harvested, washed in HEPES buffer (40 mM, pH 7.4) and adjusted to an OD_600_ of 1.0 in HEPES buffer containing either 1 mM Aldrithriol-2 (Sigma-Aldrich, Darmstadt, Germany, CAS-2127-03-9, AT-2) (oxidation control), 10 mM Dithiothreitol (Sigma-Aldrich, Darmstadt, Germany, CAS-3483-12-3, DTT) (reduction control), or buffer. As AT-2 was solved in DMSO, the solvent was added to all samples. 100 µL of the *E. coli* suspensions were added to the wells of a black, clear-bottom 96-well plate (Nunc, Rochester, NY). Fluorescence intensities were recorded using the Synergy H1 multi-detection microplate reader (Biotek, Bad Friedrichshall, Germany) at excitation wavelengths 405 and 488 nm and emission wavelength at 525 nm at 25 °C. The ratios of the fluorescence excitation intensities (405/488 nm) were used to calculate the probe’s oxidation state. All values were normalized to fully oxidized (AT-2-treated) and fully reduced (DTT-treated) roGFP2 with the following equation [1]:

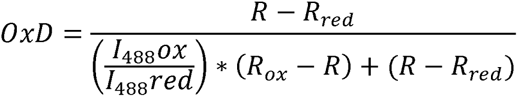

with *R_ox_* being the 405/488 ratio of oxidized (AT2-treated) and *R_red_* of reduced (DTT-treated) roGFP2 respectively. *I*_488_*ox* and *I*_488_*red* are the fluorescence intensities of roGFP2 at 488 nm under oxidizing or reducing conditions. *R* is the measured 405/488 nm ratio of roGFP2 in the respective bacterial strain under the respective condition.

**Figure 1.**
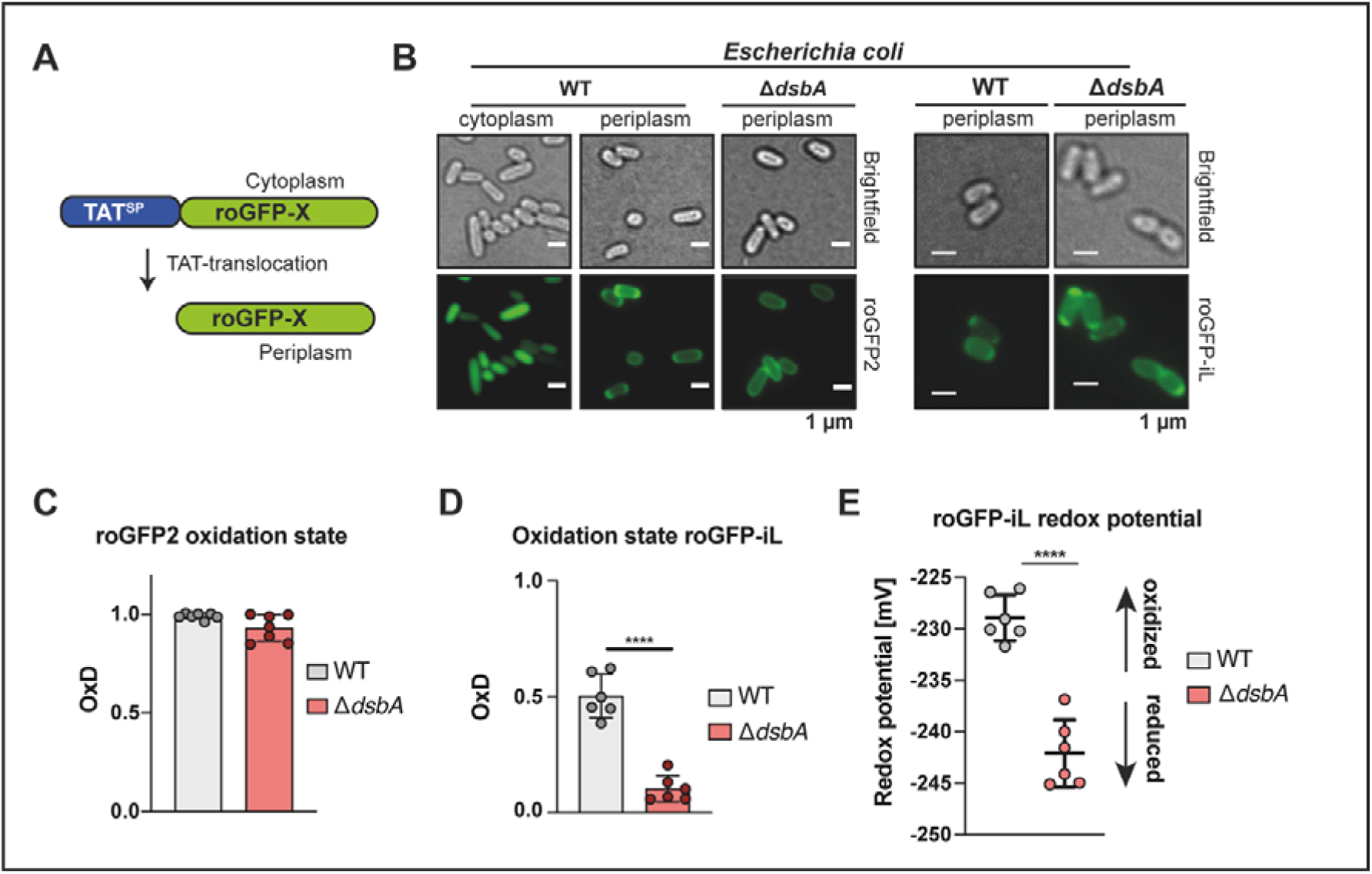
Genetically encoded redox probes reveal the redox potential imposed onto thiol pairs the periplasm of *E. coli*. **(A)** Schematic depiction of the targeting of both roGFP probes to the periplasm by N-terminal fusion with the torA signal sequence. **(B)** Correct localization of roGFP2 (left) and roGFP-iL (right) is confirmed by fluorescence microscopy. **(C)** Oxidation state (OxD) of roGFP2 in the periplasm of WT and cells lacking DsbA. The emission at 525 nm was recorded after excitation of roGFP2 at 488 nm and 405 nm in *E. coli* WT or cells lacking DsbA. OxD was determined based on samples reduced with DTT and oxidized with Aldrithiol-2 (AT-2). Each value represents the mean of three technical replicates. Error bars represent the standard deviation in n=7 individual replicates. **(D)** Oxidation state (OxD) of roGFP-iL in the periplasm of WT and of cells lacking DsbA. roGFP-iL emission at 525 nm was recorded after excitation at 395 nm or 465 nm in *E. coli* WT or Δ*dsbA*. **(E)** The roGFP-iL redox potential was calculated using the Nernst equation. Values were recorded in independent measurements and error bars represent the standard deviation in n=6 individual replicates. Significance tests in B and C were performed using Student’s t-test \*\*\*\**p*<0.0001.

For determining the re-oxidation capacity of roGFP2 in the periplasm of different *E. coli* mutants after a reductive pulse, cells expressing roGFP2 were harvested, washed in HEPES buffer and adjusted to an OD_600_ of 1.0 in HEPES buffer. Afterwards aliquots of the cell suspension were treated with 10 mM DTT for 10 min at 25°C and the reductant was washed out by three washing steps with HEPES buffer. The OD_600_ was again adjusted to 1.0 in HEPES buffer. After transfer of 100 µL of the respective cell suspensions into a black, clear-bottom 96-well plate (Nunc, Rochester, NY), 5 mM of the substances of interest: buffer, reduced (Sigma-Aldrich, CAS-70-18-8, GSH) or oxidized (Sigma-Aldrich, CAS-27025-41-8, GSSG) glutathione, cysteine or cystine, β-mercapthoethanol or DTT were added.

Fluorescence intensities were recorded as described above with 1 min 41 sec intervals for 180 min at 25 °C. Fully oxidized (AT-2-treated), fully reduced (DTT-treated) and untreated cells, that were not pre-reduced were used to calibrate roGFP2. For calculation and normalization of the probe’s oxidation degree at each time point, the means of all values recorded at all time points for *R_red_*, *R_ox_*, *I*_488_*ox* and *I*_488_*red* were used. Data was processed using Microsoft Excel software and GraphPad Prism.

### Determination of the redox potential of roGFP-iL in the periplasm of E. coli

The redox potential of roGFP-iL targeted to the periplasm of *E. coli* WT, *ΔdsbA*, Δ*gshA* or Δ*dsbA*Δ*gshA* by expression from pPT-*roGFP-iL* plasmid was determined as described previously for cytosolic roGFP2 (Degrossoli et al., 2018; Gutscher et al., 2008; Xie et al., 2020) or Grx1-roGFP2-iL located in the endoplasmic reticulum (Ugalde et al., 2022). Correct probe localization was verified by fluorescence microscopy (Supplementary figure 1). Firstly, *roGFP-iL* was expressed as described above for 16 h at 20 °C, cells were harvested, washed in HEPES buffer (40 mM, pH 7.4) and adjusted to an OD_600_ of 1.0 in HEPES buffer. The fluorescence intensity was recorded with continuous stirring every 30 sec for at least 5 min at 37 °C in an FP-8500 spectrofluorometer (Jasco, Tokyo, Japan) with a fixed emission wavelength at 510 nm. The scanned excitation wavelength ranged from 350 to 500 nm. Bandwidths of both, excitation and emission were set to 5 nm. The fluorescence excitation ratios (395/465 nm) were used as measurement of probe oxidation (Lohman and Remington, 2008). For normalization, 1 mM AT-2 and 10 mM DTT were added to the cells achieving full oxidation or reduction of the probe. For calculation, equation [1] was modified. *R_ox_* was 395/465 ratio of oxidized and *R_red_* of reduced roGFP-iL. *I*_465_*ox* and *I_465_red* represent the fluorescence intensities of roGFP2 at 465 nm under oxidizing or reducing conditions. *R* is the measured 395/465 nm ratio of roGFP-iL in the respective bacterial strain under the respective conditions. All values used for calculation of *R_ox_, R_red_, I*_465_*ox* and *I*_465_*red* are the mean of at least four measured time points, respectively.

The redox potential of roGFP-iL in the periplasm was calculated as previously described (Ugalde et al., 2022; Xie et al., 2020) using the Nernst equation [2]:

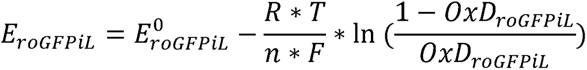

with *E^0^_roGFPiL_*=-229 mV (Lohman and Remington, 2008). *R* is the gas constant (8.314 J.K^-1^mol^-1^), *T* (310.15 K ≘ 25 °C), *n* is the number of transferred electrons (2) and *F* is Faraday’s constant (96’485 C mol^-1^). Data was processed using Microsoft Excel software and GraphPad Prism.

### Fluorescence Microscopy

*E. coli* WT, Δ*dsbA*, Δ*gshA* or Δ*dsbA*Δ*gshA* cells producing either roGFP2 or roGFP-iL from pPT plasmids cultivated as described above were harvested, washed and resuspended to an OD_600_ of 1.0 in PBS buffer (1.5 mM KH_2_PO_4_, NaCl 150 mM, 2.7 mM Na_2_HPO_4_-7xH_2_O, pH 7.4). Afterwards, 2 µL of the cell suspension were spotted on a 1.5 % Agarose pad covering a microscopy slide. *E. coli* WT cells producing roGFP2 from the pCC-roGFP2 plasmid served as cytoplasmic control. For fluorescence microscopy and image acquisition an Olympus BX51 microscope equipped with CCD camera (Retiga 3, QImaging), and LED light source (SOLA-365, Lumencor) driven by VisiView 3.× (Visitron systems) software was used. Image acquiring was carried out using a Plan-APO 100×/1.4 NA oil objective with the following filter set: BP 450–488 nm, FT 495 nm, and BP 512–542 nm. The Image J software (Schneider et al., 2012) was used for image processing.

### Intracellular roGFP2-based measurements in yeast cells

YPH499 Δ*glr1*Δ*grx1*Δ*grx2* cells were transformed with a p415TEF-*OPT1* plasmid encoding the glutathione transporter, Opt1/Hgt1, and a p416TEF plasmid encoding the indicated roGFP2 fusion construct. Cells transformed with empty plasmids served as controls for the subtraction of fluorescence background. Cells were grown at 30°C in Hartwell’s complete (HC) medium with 2% glucose and lacking leucine and uracil (-Leu-Ura) to ensure retention of both plasmids to an *OD_6_*_00_ of ∼3.0. Cells were harvested by centrifugation at 1000 x *g* for 3 min at room temperature and resuspended in 2 mL fresh HC -Ura-Leu medium containing 50 mM DTT to reduce all roGFP2 sensors before the start of the experiment. Cells were then incubated for 4 min at room temperature, centrifuged at 1000 x *g* for 3 min, washed once with 2 mL fresh HC -Ura-Leu medium and finally resuspended in fresh HC -Ura-Leu medium to an *OD*_600_ of 7.5. The cell suspension was subsequently transferred to a flat-bottomed 96-well plate, with 200 µl per well. The 96-well plate was centrifuged at 30 x *g* for 5 min so that cells formed an even, loose pellet at the bottom of each well.

To calibrate the probe, fully oxidized and fully reduced samples were generated by the addition of 20 mM *N*,*N*,*N*′,*N*′-Tetramethylazodicarboxamide (diamide, *Sigma Aldrich*) or 100 mM Dithiotreitol (DTT, *AppliChem*), respectively. Glutathione disulfide (GSSG, *Sigma Aldrich*) was added to experimental samples at final concentrations ranging from 1.5 µM to 100 µM. Probe responses were followed for 500 s in a BMG Labtech CLARIOstar^Plus^ fluorescence plate reader. All experiments were performed at least three times with cells from independent cultures. The degree of roGFP2 oxidation was calculated according to equation [1] with the exception that values were recorded at 400 and 480 nm instead of 405 and 488 nm.

### Purification of DsbAΔSP and roGFP2 from E. coli BL21 and enzymatic assay

For DsbA *in vitro* assays, DsbA was expressed as N-terminally truncated variant lacking the periplasmic signal peptide and N-terminally fused to a Strep-Tag with a TEV-cleavage site in *E. coli* BL21 cells. The protein was purified with Strep-affinity chromatography (for more details see supplementary figure 2), the tag was removed by TEV-cleavage and the protein was dialyzed into buffer W (100 mM Tris-HCl, 150 mM NaCl, 1 mM EDTA, pH 8.0). roGFP2 was expressed from pCC_*roGFP2* in *E. coli* MG1655 and purified as previously described (Müller et al., 2017). DsbAΔSP was oxidized with a 10-fold molar excess of AT-2 and roGFP2 was reduced with a 10-fold molar excess of DTT for 10 min at 20 °C and then the oxidant or reductant were removed using Micro Bio-Spin™ P-30 gel columns (BioRad, Feldkirchen, Germany, #7326223) or NAP™-5 columns (Cytiva, Freiburg, Germany, #17085302) according to the supplier’s protocol. For kinetic measurements, 0.2 µM roGFP2 were incubated with buffer alone, a 20-fold molar excess of oxidized DsbAΔSP, GSSG or both in a 96-well plate (Nunc, Rochester, NY). Addition of 2 µM AT-2 or DTT served as calibration marks for fully reduced or oxidized roGFP2 for the calculation of the OxD, according to equation [1]. roGFP2 fluorescence was recorded for 18 h at 25 °C in a Synergy H1 multi-detection microplate reader (Biotek, Bad Friedrichshall, Germany) at the excitation wavelengths 405 and 488 nm and emission wavelength at 525 nm with the following settings: 20 nm bandwidth, gain 75 %, intervals: 1 min 41 sec, optics: bottom. Data processing was carried out as described above for cellular roGFP2 assays.

**Figure 2.**
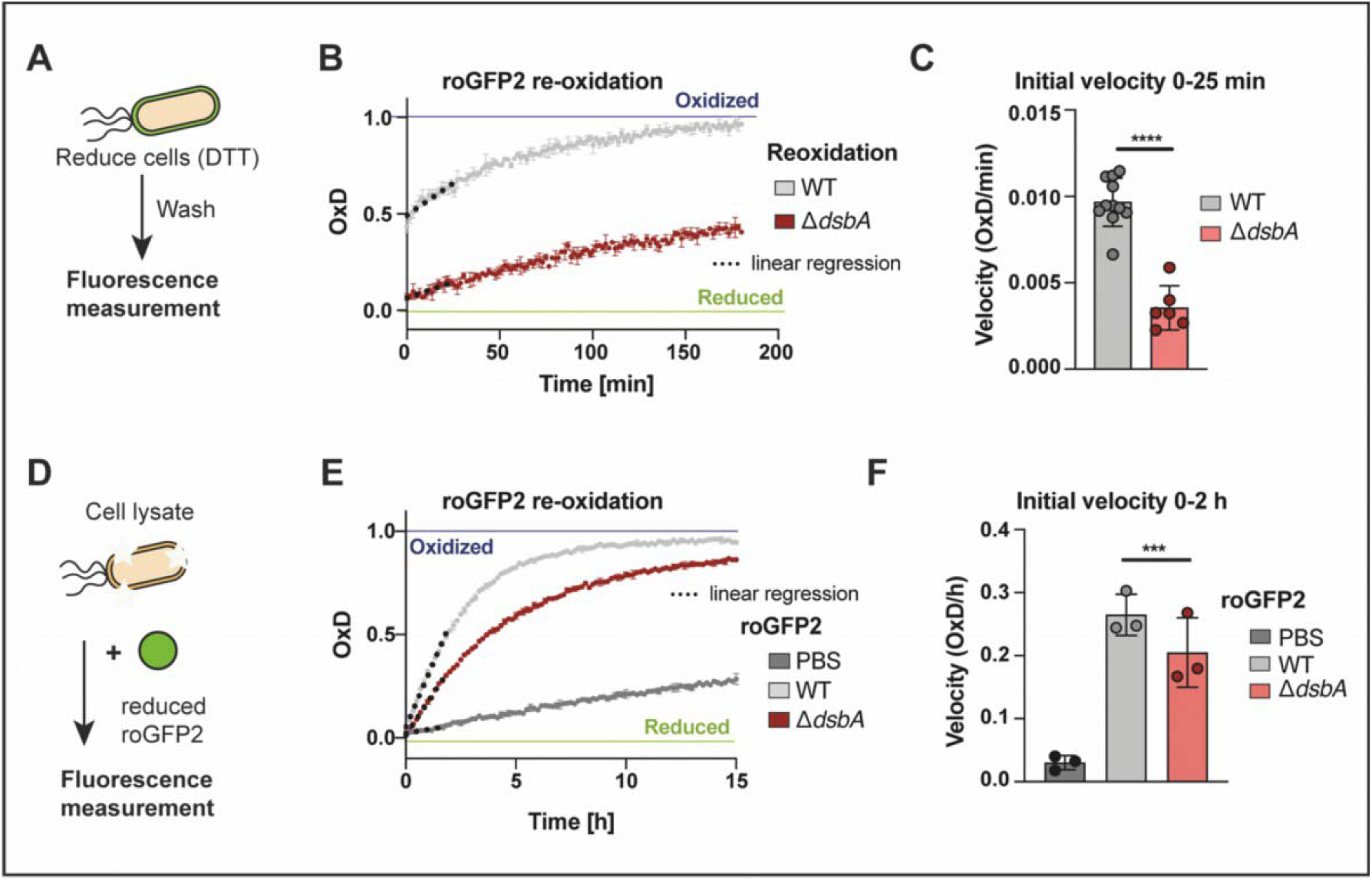
Lack of the oxidase DsbA results in inhibited re-oxidation of roGFP2 *in vivo* and *in vitro*. **(A)** Schematic representation of the workflow of *in vivo* re-oxidation kinetics in the periplasm. **(B)** Re-oxidation of roGFP2 in the periplasm of *E. coli*. *E. coli* WT and Δ*dsbA*, expressing periplasmic roGFP2 were reduced with 10 mM DTT. The reductant was washed out and fluorescence intensities were recorded. DTT-reduced and AT-2-oxidized cells served as controls for calculation of OxD. One representative example out of at least six individual replicates is shown. Error bars represent standard deviation of the technical triplicates of that individual experiment. **(C)** For an estimation of the initial re-oxidation velocity, a linear regression was calculated for the first 25 min after start of the measurement (dashed lines in B). Values shown as circles are the mean of three technical repeats recorded in at least six independent experiments. Error bars reflect the standard deviation of the means. **(D)** Schematic representation of the workflow of *in vitro* re-oxidation kinetics using *E. coli* lysate. **(E)** Oxidation of purified roGFP2 by *E. coli* lysates. Purified roGFP2 was reduced with DTT, the reductant was removed and *E. coli* cell lysates of WT or Δ*dsbA* were added (10 µg of protein). One representative experiment is shown. Values displayed are the mean of three technical replicates. Error bars reflect the standard deviation. Note the different time scales in (B) and (E). **(F)** The initial re-oxidation velocity was calculated from a linear regression in the first 2 hours from the start of the measurement (dashed lines in E). Values (circles) are the mean of three technical replicates recorded in 3 independent measurements. Error bars reflect the standard deviation of the means. Significance tests in **(C)** were performed using Student’s t-test and **(F)** using one way ANOVA. ***p< 0.001; ****p < 0.0001.

### PhoA activity assay and verification of PhoA synthesis

Activity of alkaline phosphatase PhoA was measured as described previously (Brickman and Beckwith, 1975) with modifications. Briefly, mutant strains were cultivated in 5 mL MOPS minimal medium with appropriate antibiotics at 37 °C without or with supplementation of 5 mM GSH or GSSG for around 18 h. Afterwards, *A. dest*, 5 mM GSH or GSSG were added again and the cells were incubated for one more hour. After incubation and measurement of the OD_600_, 100 mM iodoacetamide (Sigma-Aldrich, CAS-144-48-9, IAM) was added to 1 mL of the culture and cells were left on ice for 20 min. Cells were harvested (16,000 x g, 4 °C, 2 min) and washed two times with 1 mL washing buffer (50 mM NaCl, 10 mM NH_4_Cl, 10 mM MgCl_2_, 40 mM MOPS, 10 mM IAM, pH 7.3). Cell pellets were resuspended in 100 µL lysis buffer (10 mM EDTA, 2 mg/mL lysozyme, 20 mM Tris-HCl, 10 mM IAM, pH 8.0) and incubated for 30 min on ice. Afterwards, cells were lysed by three freeze and thaw cycles in liquid nitrogen and a 37 °C waterbath, followed by addition of 900 µL resuspension buffer (10 mM MgCl_2_, 10 mM ZnCl_2_, 1 M Tris-HCl, pH 8.0) and warming the samples to 28 °C. 0.04 % *p*-nitrophenylphosphate (Sigma-Aldrich, CAS-333338-18-4) were added and samples were incubated until the solution turned visibly yellow, before the reaction was stopped by adding 200 µL 1 M K_2_HPO_4_. The time point of reaction stop was noted. Lysates were then incubated on ice for 10 min and centrifuged for 5 min at 16,000 x g. The absorption at 420 nm and 550 nm of the supernatant was measured against a mixture of 100 µL lysis buffer, 900 µL resuspension buffer, 100 µL 0.4 % pNPP solution and 200 µL stop solution in a BioSpectrophotometer (Eppendorf, Hamburg, Germany). The PhoA activity was calculated using equation [3]:

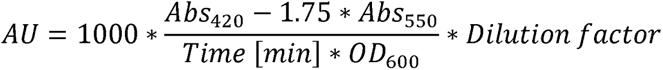

with AU being arbitrary units, Abs being the absorption at 420 nm and 550 nm and OD_600_ being the culture optical density at 600 nm.

For analysis of PhoA accumulation, 1 mL of each aforementioned cultures were harvested and the pellet was resuspended in standard 1x SDS-sample buffer adjusted to 100 µL for an OD_600_ of 1.0. PhoA accumulation was analyzed by SDS-PAGE (NuPAGE™ 4 bis 12 %, Bis-Tris, Invitrogen, Waltham, MA, USA) and Western blot analyses were carried out using nitrocellulose membranes in the iBLOT2 system (Invitrogen, Waltham, MA, USA), both according to the supplier. For detection on Western blot, a PhoA-specific first antibody (Sigma-Aldrich, #MAB1012 1:5’000) and a goat-anti-mouse IRDye 800CW conjugate (LeiCor, Lincoln, NE, USA, #926-32210) were used (1:5’000) according to the supplier’s protocol. Proteins were detected by fluorescence using the Sapphire Azure Multiimager (Azure biosystems, Dublin, CA, USA).

### Analysis of E. coli growth

*E. coli* mutant strains were cultivated in 750 µL MOPS minimal medium in a transparent 48-well plate (Sarstedt, Nümbrecht, Germany) without or with supplementation of 5 mM GSH or GSSG at 37 °C with continuous shaking in an infinite M200 multiplate reader (Tecan instruments, Männedorf, Switzerland). Cells were inoculated to a starting OD_600_ of 0.1 and bacterial growth was recorded every 30 min for 18 h.

## Results

### roGFP2 is completely oxidized in the E. coli periplasm, even in the absence of the major oxidase DsbA

The redox state of cellular compartments is often assessed by targeting genetically encoded redox probes to the compartment of interest. One frequently used redox sensor is roGFP2, a GFP variant, in which two cysteines were introduced at positions 147 and 204 that can form a disulfide bond upon oxidation. This disulfide bond results in a shift in the fluorescence excitation spectrum of the probe, in which the excitation maximum at 488 nm is reversibly decreased with a simultaneous increase of the excitation peak at 405 nm. Excitation at either maxima results in emission at ∼510 nm. The ratio of the fluorescence emission intensity upon excitation at 405 nm and 488 nm reflects changes in the dithiol disulfide state of the probe and, in combination with fully oxidized and reduced controls, allows the determination of the degree of the probe oxidation independent of probe concentration (Degrossoli et al., 2018; Dooley et al., 2004; Lukyanov and Belousov, 2014; Meyer and Dick, 2010; Xie et al., 2020). In order to assess the periplasmic oxidation state of *E. coli*, we targeted roGFP2 to this compartment. GFP is usually not folding correctly in the periplasm, hence the general secretory pathway (Sec) is unsuitable for targeting roGFP2 to the periplasm (Santini et al., 2001). In contrast, *E. coli*’s twin-arginine translocase (or Tat system) transports completely folded substrates over the inner membrane. Tat-translocation requires the presence of an N-terminal signal peptide, characterized by an essential twin-arginine motif (Palmer and Berks, 2012; Thomas et al., 2001). To target roGFP2 to the periplasm via the Tat system, we expressed the protein as an N-terminal torA_SS_ fusion in the KEIO wildtype *E. coli* BW25113 (referred to as WT) (Baba et al., 2006) (Fig. 1A). We verified translocation of roGFP2 into the periplasm using fluorescence microscopy (Fig. 1B, left panel). Determination of the roGFP2 oxidation state (OxD) revealed complete oxidation of the probe in the periplasm of *E. coli* WT (Fig. 1C, left panel). As the DsbA/DsbB pair is the major system for oxidative protein folding in the periplasm of *E. coli* (Collet and Bardwell, 2002), roGFP2 was also targeted to the periplasm of cells lacking DsbA (Fig. 1B, right panel). Surprisingly however, periplasmic roGFP2 was still fully oxidized in the absence of DsbA (Fig. 1C, right panel), suggesting the presence of an alternative mechanism for the introduction of disulfide bonds in this compartment. In both cases, due to the complete oxidation of the probe, we were not able to determine the redox potential for the roGFP2 dithiol disulfide couple.

### The redox-potential of the periplasm is significantly more reducing in cells lacking DsbA

In order to determine the periplasmic redox potential exerted on proteins containing cysteines, we turned to roGFP variants with a more oxidizing standard redox potential (Lohman and Remington, 2008). These probes, termed roGFP-iX were developed by insertion of a single amino acid (denoted by the ‘X’) adjacent to cysteine 147 resulting in a decreased stability of the disulfide bond and midpoint potentials of –229 to –246 mV compared to –280 mV of roGFP2. These roGFP-iX variants have been successfully used to monitor the oxidation state of the endoplasmic reticulum, a highly oxidized compartment in eukaryotic cells and a major site of oxidative protein folding (Delic et al., 2010; Dooley et al., 2004; Ugalde et al., 2022). Thus, roGFP-iL with a midpoint potential of –229 mV was expressed and targeted to the periplasm of the WT and Δ*dsbA* strain (Fig. 1A, B, right panel). In contrast to roGFP2, roGFP-iL was only around 50 % oxidized in the WT periplasm (Fig. 1B, left panel), allowing the calculation of the redox-potential using the Nernst equation as described before (Xie et al., 2020). In the WT the redox-potential of roGFP-iL in the periplasm was –228 mV (Fig. 1E, left panel). Not surprisingly, the lack of DsbA resulted in a more reduced roGFP-iL probe (Fig. 1E, right panel). The shift in the redox-potential was around 14 mV to –243 mV, supporting the role of DsbA in oxidative protein folding, however, the relatively small size of the shift suggests the presence of an alternative mechanism for the introduction of disulfide bonds.

### Periplasmic thiol oxidation is significantly impaired but not abrogated in a ΔdsbA-mutant

Our experiments with roGFP2 and roGFP-iL reflected the steady state of thiol oxidation in the periplasm. However, both stress and physiological situations are often accompanied by increased protein secretion, requiring correct oxidative protein folding and hence, may affect the redox homeostasis of the periplasm. In order to test the capacity of the periplasm to restore its redox balance after reductive challenge, we evaluated the re-oxidation kinetics of periplasmic roGFP2 after a reductive pulse. To explore the role of DsbA in this process, we also analyzed the re-oxidation of roGFP2 in cells lacking this oxidoreductase. In these experiments, we treated WT and cells lacking DsbA cells, both expressing roGFP2 in the periplasm with a pulse of DTT. This was followed by the removal of the reductant and the recording of the oxidation state of roGFP2 over time (Fig. 2A). In the WT, periplasmic roGFP2 oxidation starts immediately after DTT removal and restores maximum oxidation within 150 min. Cells that lack DsbA also showed sensor re-oxidation, however, roGFP2 was not restored to maximum oxidation in the time frame of our measurement (Fig. 2B).

Additionally, the re-oxidation rate, calculated as the linear slope after DTT removal, was significantly lower in DsbA-deficient cells, however, there was still re-oxidation capacity present (Fig. 2C). These findings suggest that there is an additional factor playing a role in periplasmic redox homeostasis, other than DsbA. To further explore our hypothesis, we used cell lysates derived from *E. coli* WT and DsbA-deficient cells and explored their capacity to re-oxidize DTT-reduced, purified roGFP2 (Fig. 2D). Similar to the cell re-oxidation assay, we calculated the oxidation rate from the linear slope within the first two hours of measurement. Although roGFP2 oxidation by cell lysate was slow, when compared to intact cells, lysate from WT was significantly faster in catalyzing roGFP2 re-oxidation than a Δ*dsbA* lysate.

However, in line with our previous findings, indicating another factor capable of catalyzing disulfide bond formation in the periplasm, the Δ*dsbA* lysate was still significantly faster than a buffer control (Fig. 2E, F). The rather slow oxidation of roGFP2 *in vitro* may be explained by the fact that DsbA preferably introduces disulfide bonds in unfolded proteins entering the periplasm over folded ones (Kadokura et al., 2004).

### Oxidized glutathione rescues periplasmic roGFP2 re-oxidation velocity in cells lacking DsbA

Based on our observations, we suspected that glutathione (GSH) and its oxidized dimer GSSG might influence the periplasmic redox balance in *E. coli* and thus might act as part of an auxiliary system in oxidative folding. Glutathione is one of the most important redox-active molecules inside many cells and it has been shown to be present in the periplasm (Eser et al., 2009; Pittman et al., 2005).

We thus analyzed the periplasmic capacity for re-oxidizing roGFP2 in the presence of glutathione after a reductive pulse. *E. coli* WT and DsbA-deficient cells producing periplasmic roGFP2 were thus reduced with DTT, and, after reductant removal, roGFP2 oxidation was recorded in the presence of 5 mM GSH or GSSG (Fig. 3A). Surprisingly, addition of GSH to WT cells resulted in an accelerated roGFP2 re-oxidation in the periplasm, whereas GSSG supplementation had no discernible influence (Fig. 3B). In contrast, in cells that lack DsbA, GSH and GSSG acted in a more expected way; while GSSG supplementation rescued roGFP2 re-oxidation, GSH had no influence on the periplasmic redox dynamics in Δ*dsbA* (Fig. 3C). Calculation of the re-oxidation rate confirmed that while GSH significantly speeds up re-oxidation of periplasmic roGFP2 in the WT, GSSG significantly accelerates the re-oxidation rate of roGFP2 in DsbA-deficient cells, essentially to WT level (Fig. 3D). Based on these findings, we asked ourselves if DsbA is interacting with glutathione, causing the observed opposite responses to GSH and GSSG in WT and DsbA-deficient *E. coli* cells.

**Figure 3.**
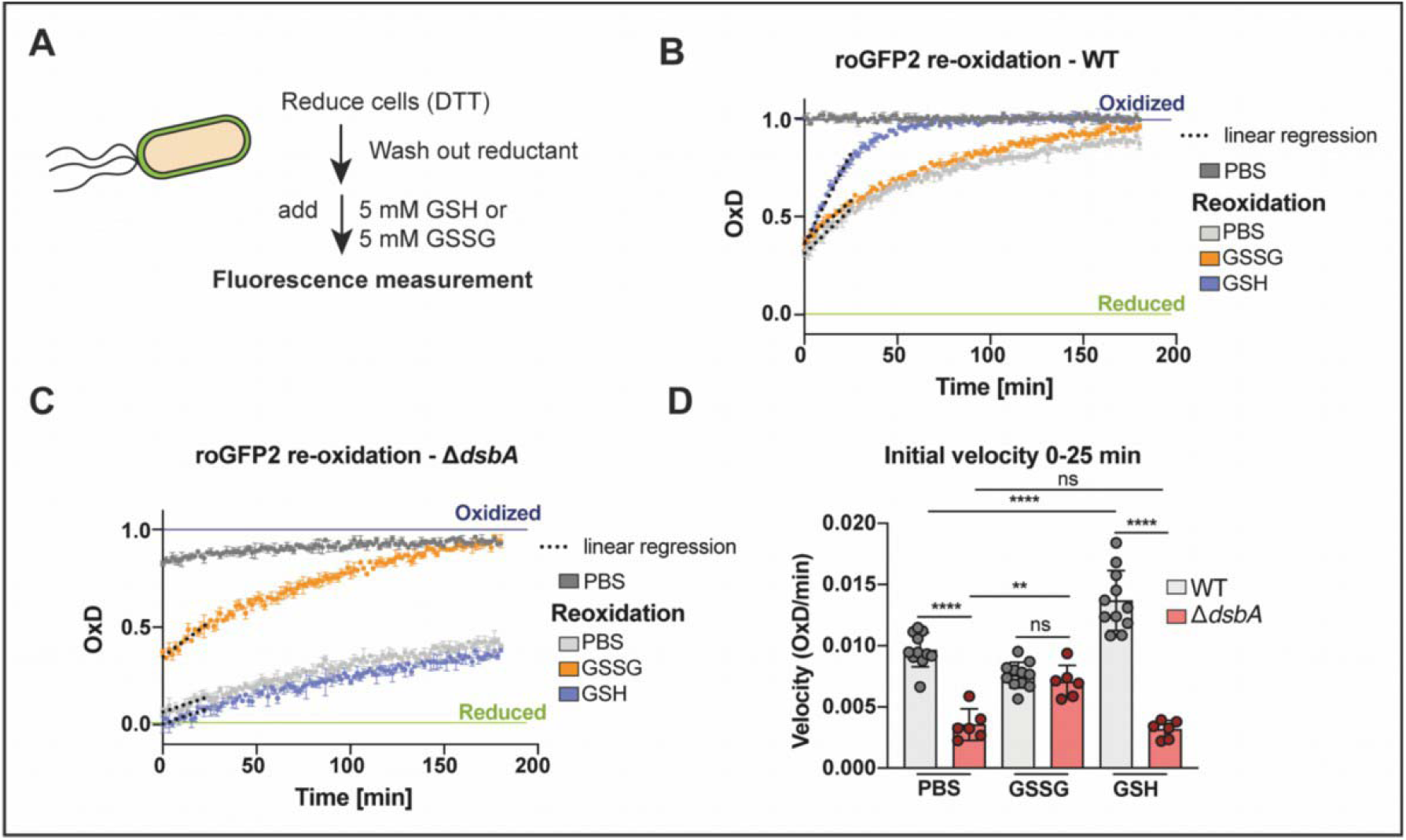
External addition of oxidized glutathione rescues the re-oxidation of roGFP2 in the periplasm of cells lacking DsbA, while reduced glutathione accelerates re-oxidation of roGFP2 in the periplasm of *E. coli* WT. (**A**) Schematic representation of the workflow of *in vivo* re-oxidation kinetics in the periplasm. *E. coli* WT (**B**) and Δ*dsbA* (**C**) expressing periplasmic roGFP2 were reduced with 10 mM DTT. The reductant was washed out, 5 mM GSH or GSSG were added and fluorescence intensities were recorded. DTT-reduced and AT-2-oxidized cells served as controls for calculation of OxD. A representative experiment is shown. Values are the mean of three technical replicates. Error bars reflect the standard deviation. (**D**) The initial re-oxidation velocity was calculated as linear regression for the first 25 min after start of the measurement (dashed lines in B and C). Values (circles) are the mean of three technical replicates recorded in a minimum of six experimentally independent replicates. Error bars represent the standard deviation. Values for PBS treated cells of the WT and Δ*dsbA* are the same as shown in Fig. 2 and are presented here for context. Significance test was performed using one way ANOVA. ns>0.05, \*\**p*<0.01, \*\*\*\**p* < 0.0001.

### Oxidized glutathione does not directly interact with the oxidase DsbA

To investigate if glutathione interacts with DsbA, we investigated the effect of GSSG on DsbA-dependent roGFP2 oxidation *in vitro*. Purified DsbA was oxidized and added to reduced roGFP2. Similar to the aforementioned lysate assay, the oxidation rate of roGFP2 by DsbA is rather slow compared to *in vivo* re-oxidation, although significantly higher compared to buffer alone (Fig. 4A, B). The oxidation rate of roGFP2 in the presence of GSSG alone was slightly slower, but still comparable to roGFP2 oxidation by DsbA. Adding both GSSG and DsbA at the same time did not even double the probe’s oxidation rate, indicating an additive effect of GSSG and DsbA and not a GSSG-driven catalytic action of DsbA.

**Figure 4.**
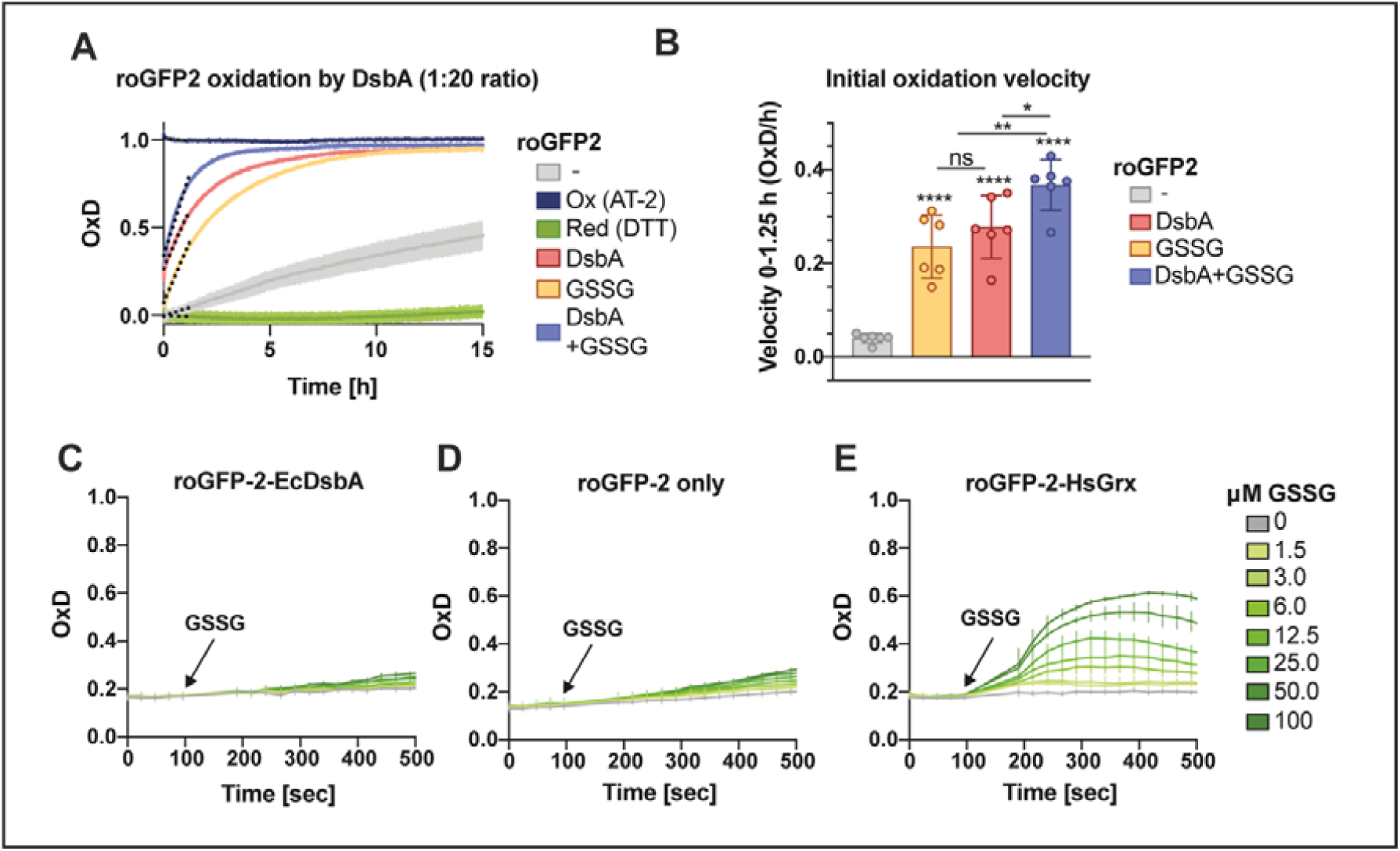
Oxidation of roGFP2 by oxidized glutathione (GSSG) is not catalyzed by DsbA. **(A)** Oxidation of purified reduced roGFP2 by AT-2-oxidized DsbA (red), GSSG (orange) or both (purple). Purified roGFP2 was reduced with DTT, the reductant was washed out and DsbA and/or GSSG were added in a 20-fold molar access. PBS was used to track air oxidation of roGFP2. DTT served as reduction and AT-2 as oxidation controls for calculation of OxD. **(B)** The initial re-oxidation velocity was calculated as linear regression for the first 1.25 hours after the start of measurement (dashed lines A). Values (circles) are the mean of three technical replicates recorded in a minimum of six independent repeats and error bars depict standard deviation. Significance test was performed using one-way ANOVA ns *p*>0.05 *\*p*<0.05, \*\**p*<0.01, \*\*\**p*<0.001, \*\*\*\**p* < 0.0001. **(C)** The oxidation state of the cytosolic roGFP2 fusion protein roGFP2-*Ec*DsbA was monitored in ScOpt1/ScHgt1 YPH449 *glr1 grx1 grx2* yeast cells upon exogenous addition of different GSSG amounts. Unfused roGFP2 **(D)** served as negative and roGFP2-*Hs*Grx1 **(E)** as positive control for protein-GSSG interaction. Values **(B-E)** are the mean of four individual replicates and error bars depict the standard deviation.

To further confirm the incapability of DsbA to perform GSSG-dependent roGFP2 oxidation, we used the cytosol of genetically manipulated yeast cells as “cellular test tubes”. This assay is performed in yeast cells, lacking the glutathione reductase (Glr1) and both cytosolic class I dithiol glutaredoxins (Grx1, Grx2), while simultaneously expressing Opt1, a glutathione transporter (Zimmermann et al., 2021). This *in vivo* system should enable us to monitor a potential DsbA-catalyzed oxidation of roGFP2 by GSSG in the absence of any *E. coli*-specific factors. For this, we engineered a roGFP2-DsbA fusion construct and expressed it in the cytosol of the Δ*glr*Δ*grx*1Δ*grx*2 yeast cells, following the exogenous application of GSSG concentrations between 1.5-100 µM. Unfused roGFP2 served as negative and roGFP2 fused to *homo sapiens* glutaredoxin (roGFP2-*Hs*Grx) as positive control for direct interaction with GSSG. In this assay, GSSG-driven roGFP2 oxidation did not depend on the oxidase DsbA (Fig. 4C, D). However, we cannot exclude that roGFP2 is an inappropriate substrate of DsbA in the fusion construct and therefore roGFP2 may not be oxidized by DsbA in this assay. In contrast, the roGFP2-*Hs*Grx fusion probe strongly reacted to the addition of GSSG (Fig. 4E). Both the *in vitro* and the “cellular test tube” approach strongly suggest that DsbA does not interact with glutathione itself, indicating that the role of glutathione in the periplasmic redox homeostasis is independent of the known mechanism for disulfide bond formation in *E. coli*.

### Monothiols accelerate roGFP2 re-oxidation in a DsbA-dependent mechanism

While we excluded the interaction of GSSG with DsbA, we showed that the addition of GSH accelerated the re-oxidation rate of periplasmic roGFP2 after a reductive pulse in a DsbA-dependent manner (Fig. 3B, D). To investigate whether this effect is limited to GSH, we tested the influence of other reduced monothiols and dithiols. Cystein, β-mercaptoethanol, and DTT were added to *E. coli* WT and Δ*dsbA* cells and roGFP2 re-oxidation dynamics in the periplasm were measured as described above. The addition of the monothiols cysteine and β-mercaptoethanol to WT significantly accelerated roGFP2 oxidation similar to GSH, as measured by the time after which the probe reached full oxidation. In contrast to monothiols, addition of the dithiol DTT to WT completely inhibited roGFP2 re-oxidation. We also included cystine in our assay, the oxidized form of the amino acid cysteine. In contrast to GSSG, which had no impact on the roGFP2 oxidation rate in WT, cystine massively reduced the time until full oxidation was reached (Fig. 5A, B). In the Δ*dsbA* mutant, the presence of monothiols had no significant impact, while cystine supplementation caused a drastically increased re-oxidation state and rate, similar to WT (Fig. 5C, D). Overall, these findings suggest a DsbA-dependent effect of monothiols, accelerating thiol oxidation in the periplasm, even though we did not observe direct interaction of oxidized glutathione with DsbA itself.

**Figure 5.**
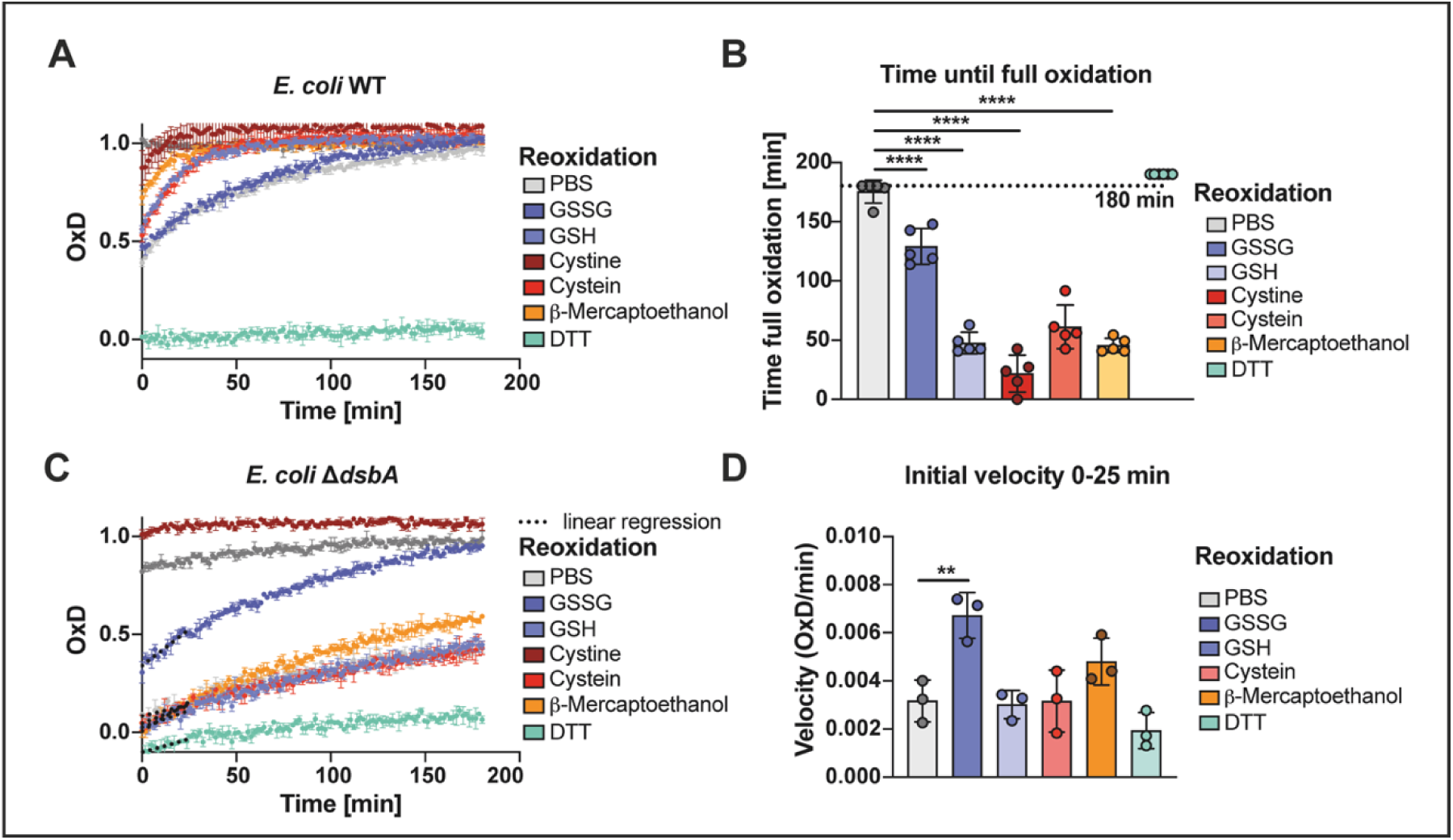
The re-oxidation of periplasmic roGFP2 in *E. coli* WT is accelerated by external addition of different monothiols. **(A)** Influence of monothiols on the re-oxidation of roGFP2 in *E. coli* periplasm after a reductive pulse. *E. coli* WT cells with roGFP2 in their periplasms were reduced with 10 mM DTT, before the reductant was washed out and 5 mM GSH, GSSG, cysteine, cystine, DTT or β-mercaptoethanol were added. Then, fluorescence intensities were recorded for 3 h. DTT-reduced and AT-2-oxidized cells served as controls for calculation of OxD. Values displayed are the mean of three technical replicates representative for five independent experiments. Error bars reflect the standard deviation. **(B)** The time after which roGFP2 was completely oxidized was calculated to visualize differences in the re-oxidation speed. Values (circles) are the mean of three technical replicates recorded in five independent measurements. DTT-treatment did not result in re-oxidation during the 180 min measurement time. Error bars reflect the standard deviation of the means. Significance test was performed using one way ANOVA. \*\*\*\**p* < 0.0001. **(C)** Influence of monothiols on the re-oxidation of roGFP2 in the periplasm of *E. coli* cells lacking DsbA after a reductive pulse. Values shown are the mean of three technical replicates, representative for three independent experiments. Error bars depict the standard deviation. **(D)** The initial re-oxidation velocity was calculated as linear regression for the first 25 min after start of the measurement (dashed lines A). Values (circles) are the mean of three technical replicates recorded in three independent repeats and error bars depict the standard deviation of those means. Significance test in was performed using one-way ANOVA ns *p*>0.05, \*\**p*<0.01.

### Endogenous glutathione is involved in stabilizing and maintaining the periplasmic redox state

All our experiments thus far were performed with exogenous glutathione. But we also wondered about the role of endogenous glutathione, synthesized in *E. coli*’s cytoplasm. For this, we determined the oxidation state of periplasmic roGFP-iL in cells lacking GshA, the first enzyme of *E. coli*’s glutathione biosynthesis pathway. In glutathione-free media, these cells do not contain GSH (Apontoweil and Berends, 1975; Carmel-Harel and Storz, 2000). Confirming our observation with exogenous GSH, our experiment revealed a slightly, but significantly lower oxidation of roGFP-iL in the periplasm of cells lacking GSH. The redox potential shifted from around –228 mV (WT) to –233 mV, counterintuitively a more reducing state, in the absence of GSH (Fig. 6A, B). Nevertheless, the shift was not as pronounced as in cells lacking DsbA, which had a roGFP-iL redox potential of around –243 mV (Fig. 1E).

**Figure 6.**
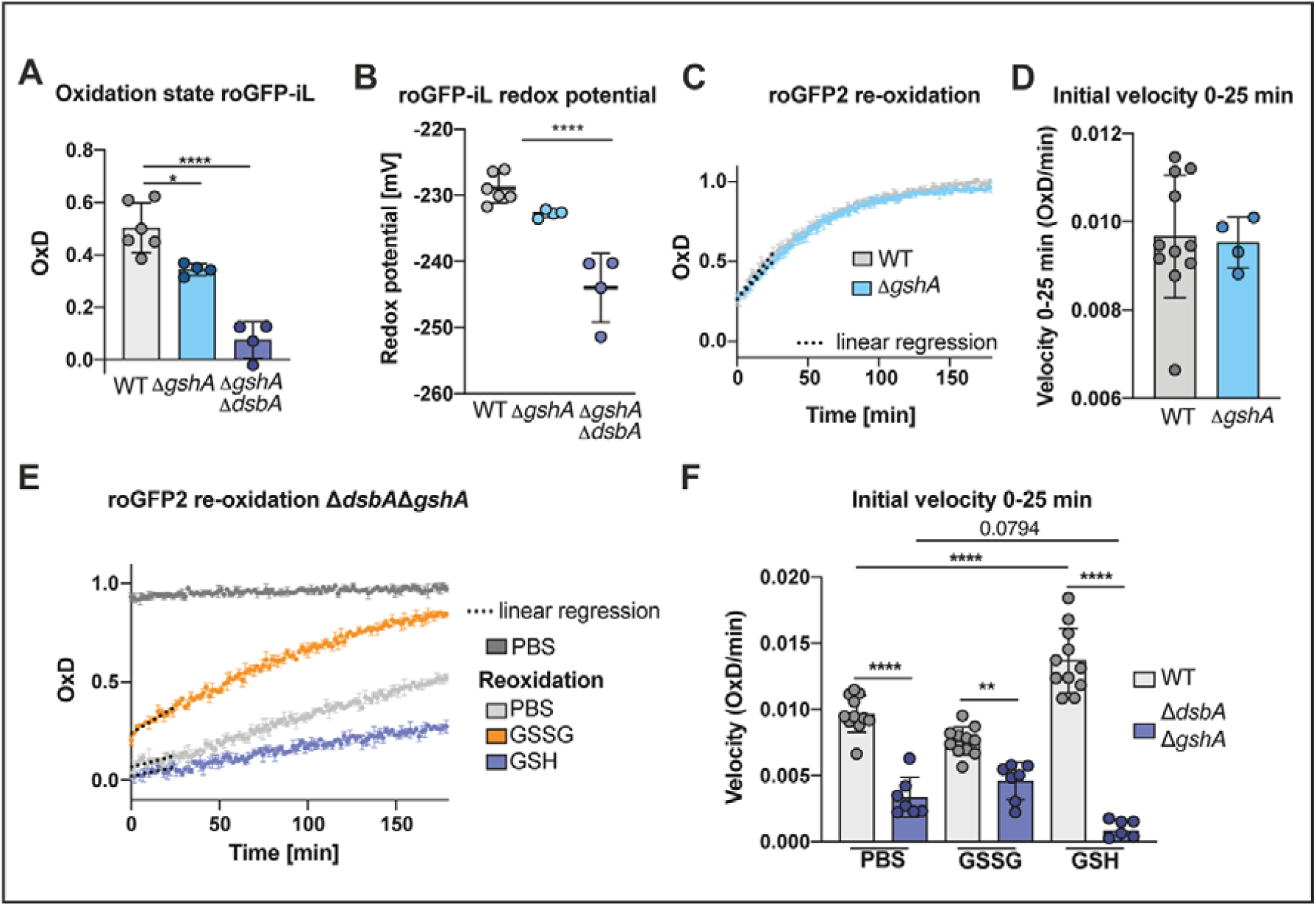
Expression of roGFP probes in cells lacking DsbA and analysis of the periplasmic redox potential and re-oxidation rates indicate that endogenous glutathione is involved in stabilizing and maintaining the periplasmic redox state. (**A**) Oxidation state (OxD) of roGFP-iL in the periplasm of WT and cells lacking GSH. The emission at 525 nm was recorded after excitation at 395 nm or 465 nm of roGFP-iL in the respective cells. Oxidized or reduced roGFP-iL was generated by treatment of cells with AT-2 or DTT. (**B**) The periplasmic redox potential was calculated from (A) using the Nernst equation. The Values (circles) are the mean of three technical replicates recorded in a minimum of four independent repeats and error bars depict standard deviation of the means. Values for the WT were already shown in Fig. 2 and are presented again for context. Significance tests in A and B were performed using ANOVA-test. \*\*\*\**p*<0.0001. (**C**) Re-oxidation of roGFP2 in the periplasm of *E. coli* WT and Δ*gshA*. The assay was performed as described in Fig. 2. One representative example out of at least four individual repeats is shown. Error bars represent standard deviation of technical triplicates. (**D**) The initial re-oxidation velocity was calculated from linear regression for the first 25 min after start of the measurement (dashed lines in B). Values shown as circles are the mean of three technical repeats recorded in at least four independent assays. (**E**) Re-oxidation of periplasmic roGFP2 in *E. coli* Δ*gshA*Δ*dsbA*. One representative example out of at least seven individual repeats is shown. Error bars represent standard deviation of technical triplicates. (**F**) The initial re-oxidation velocity was calculated from linear regression in the first 25 min after start of the measurement (dashed lines). Values (circles) are the mean of three technical replicates recorded in a minimum of seven independent repeats. Error bars represent the standard deviation. Significance test was performed using one way ANOVA. \*\**p*<0.01, \*\*\*\**p* < 0.0001. Values for the WT were already shown in Fig. 3 and are presented again for context.

We also asked whether the presence of endogenous GSH leads to faster re-oxidation of roGFP2. To address this question, we analyzed the capacity to restore the redox balance after reductive challenge in GSH-depleted cells producing periplasmic roGFP2 as described before. Our data indicates that roGFP2 oxidation rate and end oxidation state in the periplasm of GSH-deficient cells was comparable to WT (Fig. 6C, D) suggesting GSH is not essential for recovery after a reductive challenge.

Next, we asked what happens when both, DsbA and GSH are missing. The redox state of roGFP-iL in a Δ*gshA*Δ*dsbA* strain showed a periplasmic redox potential imposed on the sensor (ca. –244 mV) similar to the redox potential in the *dsbA* single mutant (ca. –243 mV) (Fig. 1, Fig. 6B). Intriguingly however, using a Δ*gshA*Δ*dsbA* strain in a periplasmic roGFP2 re-oxidation assay revealed that exogenous GSSG did not completely restore WT oxidation rate and final oxidation state (Fig. 6E, F) contrary to cells lacking solely DsbA (Fig. 3).

Adding GSH to a Δ*gshA*Δ*dsbA* strain even decelerated periplasmic roGFP2 oxidation, something we did not observe in the Δ*dsbA* single mutant (Fig 3). These experiments indicate that a fine-tuned balance of reduced and oxidized glutathione plays a role in stabilizing and maintaining the redox environment in the periplasm of *E. coli*, and cells lacking both DsbA and endogenous glutathione can neither fully utilize the oxidative power of exogenous GSSG nor compensate the reductive power of GSH in their periplasm.

### Endogenous glutathione is involved in disulfide bond formation in E. coli’s own periplasmic proteins

The observed influence of GSH on growth phenotypes (Supplementary figure 3) suggests that our previous observations of disulfide bond formation in heterologously expressed roGFP-based redox sensors also apply to endogenous DsbA substrates. One well-characterized substrate of DsbA is alkaline phosphatase PhoA. PhoA is only active upon formation of intramolecular disulfide bonds essential for correct protein folding. The activity of this enzyme can be measured in a colorimetric assay using the substrate *para*-nitrophenolphosphate (Fig. 7A) (Berg, 1981; Brickman and Beckwith, 1975; Sone et al., 1997). We thus tested the activity of alkaline phosphatase PhoA in different *E. coli* deletion strains grown in MOPS minimal medium. As described above, the lack of GSH, although without effect on re-oxidation of roGFP2 in the periplasm, slightly shifted the periplasmic redox homeostasis to more reducing conditions, as seen in the lowered redox potential of roGFP-iL in those cells (Fig. 6). In accordance with this, PhoA activity was slightly, but significantly lowered in GSH-deficient cells compared to WT. Also, in line with our previous results (Fig. 1), cells lacking DsbA showed only poor PhoA activity, and the same was true for cells lacking both DsbA and GSH (Fig. 7B).

**Figure 7.**
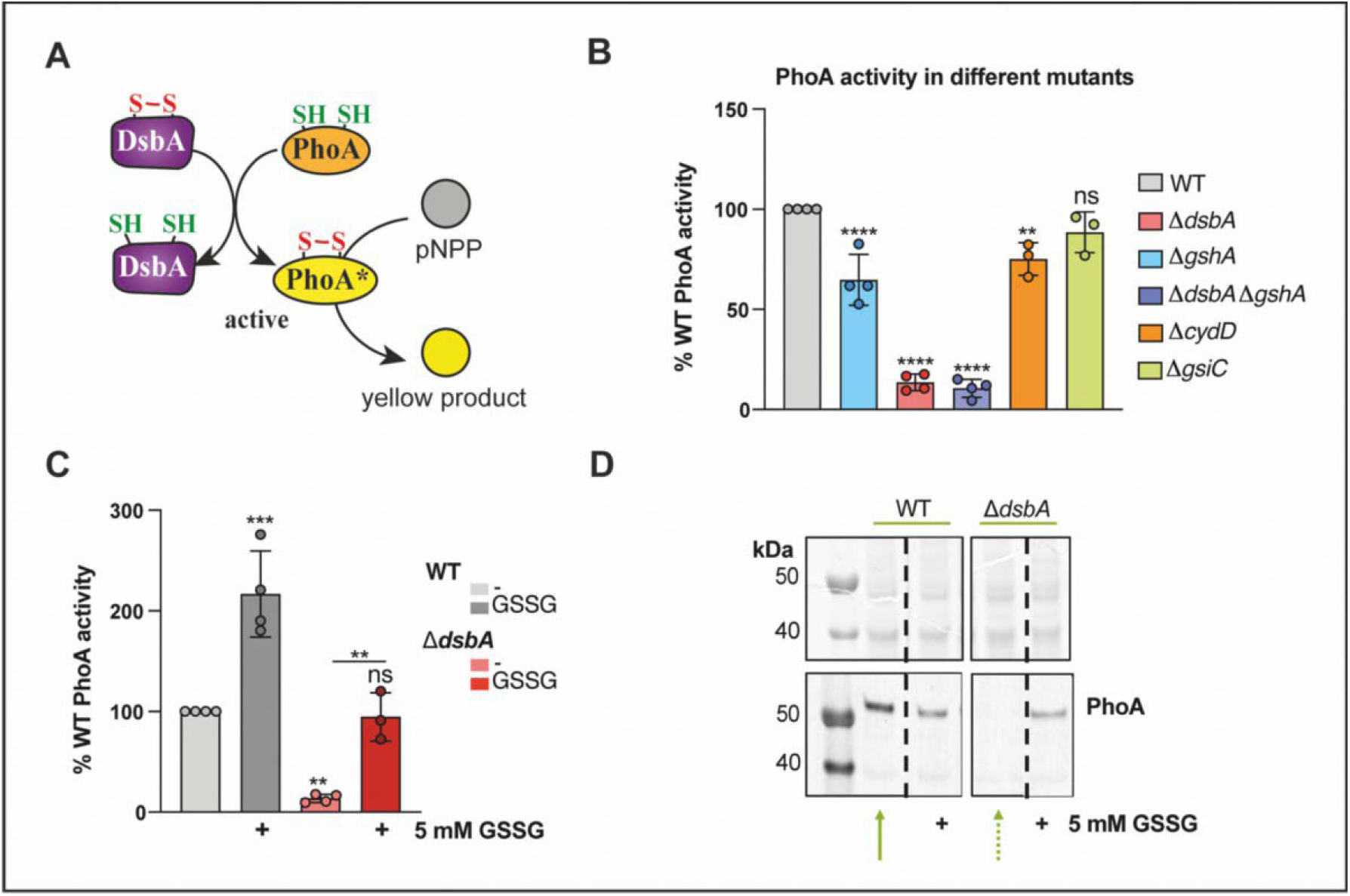
Endogenous alkaline phosphatase (PhoA) is less active in the absence of glutathione and DsbA in the periplasm, and GSSG restores oxidative folding of PhoA in the absence of DsbA. (**A**) PhoA activity depends on oxidation by DsbA and activity of PhoA can be monitored using *p*-nitrophenylphosphate (pNPP) resulting in formation of the yellow product *p-*nitrophenol. (**B**) PhoA activity was assayed in cells cultivated in MOPS medium for 24 h at 37 °C. Release of *p*-nitrophenol from the PhoA substrate pNPP was measured and relative PhoA activity in % of WT activity was calculated. (**C**) GSSG restores PhoA activity to WT level in cells lacking DsbA. Cells were cultivated in MOPS medium in the presence of 5 mM GSSG for 18 h at 37 °C. Then fresh GSSG (5 mM) was added and the cells were incubated for 1 h. PhoA activity was assayed as described above. (**D**) PhoA accumulation was analyzed by SDS-PAGE and western blot in cells cultivated as described in (C). For western blot a PhoA specific first and a fluorophore-coupled second antibody were used and fluorescence signal was scanned. Values in B and C are the mean of at least three independent replicates of duplicate assays with error bars representing the standard deviation. Significance test was performed using one way ANOVA. ns *p*>0.05 *\*p*<0.05, \*\**p*<0.01, \*\*\**p*<0.001, \*\*\*\**p* < 0.0001.

To test whether the observed reduction in PhoA activity is dependent on glutathione levels in the periplasm, we also measured PhoA activity in a *cydD* mutant. This mutant can still synthesize glutathione, but is not able to produce the inner membrane ABC-transporter CydDC that transports glutathione from the cytosol into the periplasm and hence suffers from reduced periplasmic glutathione levels (Mironov et al., 2020; Pittman et al., 2005; Shepherd, 2015). Again, we observed a slight, but significant reduction of PhoA activity in this strain, although less pronounced than in cells completely lacking glutathione biosynthesis (Fig. 7B). Conversely, blocking glutathione transport from the periplasm into the cytosol by deleting GsiC, the inner membrane component of the GsiA-D ABC Transporter (Suzuki et al., 2005; Wang et al., 2017, 2018) did not result in significant changes in PhoA activity compared to the WT.

Next, we supplemented the growth medium with exogenous GSSG to test if, in accordance with the re-oxidation assays (Fig. 3) and the rescue of the growth phenotype (Supplementary figure 3), it can assist in oxidative folding of PhoA in the absence of DsbA. Indeed, GSSG addition restored PhoA activity back to WT level (Fig. 7C). Furthermore, addition of exogenous GSSG also increased PhoA activity in the WT, contrary to what we observed with roGFP2, suggesting that the role of periplasmic glutathione in oxidative folding differs, depending on the protein in question (Fig. 3, Supplementary figure 3).

Proteins that are not correctly folded are usually unstable in the periplasm (Hiniker and Bardwell, 2004), and thus no PhoA protein could be detected on a western blot in DsbA-deficient cells (Fig. 7D). External GSSG addition prevented PhoA from degradation in a Δ*dsbA* mutant.

Taken together, our data indicates a significant role for glutathione, not only in oxidative protein folding of periplasmic proteins, but also by balancing and maintaining the periplasmic redox homeostasis.

## Discussion

The DsbA/B system is the major thiol oxidation system in the periplasm of *E. coli* and together with the DsbC/D thiol disulfide isomerase system forms the oxidative folding machinery (Collet and Bardwell, 2002; Manta et al., 2019). The standard redox potentials of those systems are known (Wunderlich et al., 1993; Zapun et al., 1995), but the *in vivo* redox potential seen by protein thiol disulfide pairs in the periplasm is unknown. Here, we used the genetically encoded proteins roGFP2 and roGFP-iL targeted to the periplasm to directly probe the thiol redox homeostasis in this compartment. When expressed in the cytosol of *E. coli*, the probe roGFP2 with a midpoint redox potential of –280 mV is almost completely reduced (Degrossoli et al., 2018). However, we found that it is virtually fully oxidized, when expressed in the periplasm, suggesting that its engineered cysteine residues are completely oxidized by the oxidative folding machinery.

To our surprise, roGFP2 was also fully oxidized in cells lacking the major thiol oxidase in the periplasm, DsbA. We thus also analyzed the re-oxidation capacity after a reductive pulse, as the oxidation state in an unperturbed cell solely reflects the steady state. And in the absence of DsbA we did indeed find a significantly diminished re-oxidation velocity underlining the importance of DsbA for oxidative folding.

Using roGFP-iL, we were able to determine that the redox potential imposed onto a thiol pair in the periplasm is -229 mV. In the absence of DsbA, this redox potential shifted to a more reducing -243 mV. In a previous study, Messens et al. used an Ag/AgCl electrode to measure the redox potential in *E. coli* WT periplasmic extracts (Messens et al., 2007). Typically, an electrode will measure the redox potential of all species it can chemically interact with and in this case the redox potential was determined to be -165 mV. Unexpectedly, in their setting, removal of DsbA shifted the redox potential to an even more oxidizing redox potential. Our finding that the thiol disulfide redox potential is significantly below the overall redox potential observed by Messens et al. with an electrode could explain their seemingly paradoxical finding, since in a Δ*dsbA* strain a component that is reducing in comparison to the overall redox potential measured by the electrode is removed from the overall redox pool. In our case, removal of DsbA did indeed result in an overall more reducing redox potential imposed on thiol pairs.

In concordance with our results in the periplasm, roGFP2 targeted to the ER is fully oxidized, as well. Similarly, a roGFP2-iL-Grx1 fusion redox probe targeted to the ER of *Arabidopsis thaliana* cells lacking Ero1/2, the functional homolog of DsbB in eukaryotes was more reduced, and re-oxidation after a reductive pulse was inhibited by the lack of Ero1/2 (Ugalde et al., 2022).

While roGFP2 re-oxidation was diminished in Δ*dsbA*, it was not absent, suggesting to us the presence of an alternative pathway for disulfide bond formation. As glutathione is one of the major redox buffers in cells, we assumed a possible role for the small molecule in periplasmic redox homeostasis. Glutathione’s cytosolic functions have been studied extensively and, until a few years back, GSH was thought to be absent from the periplasm (Pittman et al., 2005). However, relatively high glutathione levels in the periplasm of *E. coli* were discovered recently, and since, there is an ongoing debate on its function in this compartment (Delaunay-Moisan et al., 2017; Eser et al., 2009; Pittman et al., 2005). Interestingly, cells lacking the transporter for GSH from the cytosol to the periplasm, CydDC, exhibit several phenotypes, including DTT sensitivity and swarming defects, also found in cells deficient in oxidative protein folding. This is an additional indication for a role of GSH in redox homeostasis and oxidative folding in the periplasm. (Fabianek et al., 2000; Goldman et al., 1996; Pittman et al., 2005; Smirnova et al., 2012). Accordingly, we found that re-oxidation of periplasmic roGFP2 in Δ*dsbA* was restored by the addition of exogenous GSSG, while it was not influenced in WT. Surprisingly, adding exogenous GSH to the WT accelerated roGFP2 re-oxidation, but not in a Δ*dsbA* strain. We also observed this seemingly paradoxical acceleration of thiol re-oxidation in WT by other reducing monothiols like cysteine or β-mercaptoethanol. However, we did not observe direct interactions of DsbA with glutathione *in vitro* and in our yeast cell experiments.

We next assessed the role of endogenous glutathione in the redox balance of the periplasm. And in line with our observations with exogenous GSH, the oxidation state of roGFP-iL was slightly shifted to a more reducing state in Δ*gshA*, indicating a role for endogenous glutathione in the periplasmic redox homeostasis as well. It should be noted, however, that roGFP2 re-oxidation in a GSH-deficient mutant was comparable to WT. Taken together, the presence of GSH and GSSG in the periplasm, driven by the availability of exogenous and endogenous glutathione, seems to be important for the fine-tuning of the periplasmic redox potential.

Expression of a non-native redox sensor might not reflect the natural redox state in the periplasm or might even influence it by diverting oxidative power available for the formation of native disulfide bonds. We thus analyzed the activity of PhoA, a native *E. coli* protein, which depends on oxidation by DsbA for its activity, in different mutants. This approach revealed that reduced periplasmic glutathione levels indeed resulted in significantly lower PhoA activity, in line with the more reduced roGFP-iL redox state in Δ*gshA*. While PhoA’s oxidative folding is clearly influenced by the presence or absence of GSH, not all DsbA substrates seem to be influenced by glutathione. RNaseI folding and isomerization by DsbA/DsbC e.g., was not influenced by the loss of GSH (Messens et al., 2007).

In order to understand the role of glutathione in the periplasm, it is helpful to have a look at the role of GSH in the eukaryotic ER, for which, in contrast to bacteria, more studies are available. In the ER, the glutathione concentration is around 15 mM, higher than in whole cell lysates with around 7 mM (Birk et al., 2013). Similar to the periplasm, the GSH:GSSG ratio in the ER is lower (3:1 to 1:1) compared to the cytosol (100:1), resulting in a more oxidizing environment (Hwang et al., 1992). It is discussed whether GSH is the reductive power for disulfide isomerization by PDI (Vitu et al., 2010) and high GSSG levels could act as oxidant reservoir; however, it is still unclear if GSSG itself is able to oxidize PDI (Lappi and Ruddock, 2011; Ushioda and Nagata, 2019). In the periplasm of Gram-negative bacteria, DsbB recycles DsbA, but in contrast to Ero1, it uses the respiratory chain as electron sink. As aforementioned, we observed accelerated roGFP2 re-oxidation in the WT, but not in Δ*dsbA*, when adding monothiols to the cells and the opposite effect for GSSG, compensating the lack of DsbA, raising the question whether DsbB is somehow regulated by glutathione. However, analyzing a *dsbB* mutant strain regarding its oxidation state and re-oxidation capacity in presence or absence of GSH or GSSG revealed that GSSG was still able to complement for the loss of DsbB, indicating a DsbB-independent mechanism (Supplementary figure 4).

Overall, we showed that oxidized glutathione can compensate for the loss of DsbA by an unknown mechanism. One possibility is that oxidized glutathione directly oxidizes roGFP2 and other reduced proteins, however previous studies (Müller et al., 2017) and the current *in vitro* and yeast data show that direct roGFP2 oxidation by glutathione is very inefficient.

Another possibility is the presence of a yet unidentified redox factor in the periplasm that can catalyze oxidative protein folding in the absence of DsbA. One possibility is DsbC and it has been suggested that reduced glutathione can react with the isomerase, especially by providing reductive power when DsbD is missing (Pittman et al., 2005; Smirnova et al., 2012).

However, (Messens et al., 2007) could also show that DsbC alone was not able to substitute for DsbA in folding of RNaseI. In the ER up to 20 different oxidoreductases, for example the peroxiredoxin Prx4 or the glutathione peroxidase-like enzymes Gpx7 and Gpx8 are found besides PDI and for at least some of them it has been proposed that they possibly oxidize PDI (Nguyen et al., 2011; Wang et al., 2018; Zito et al., 2010). We think it is possible, that *E. coli* has a similar backup system for DsbA, presumably a glutaredoxin-like protein, which is coupled to the periplasm’s glutathione pool, providing either oxidizing or reducing power.

Taken together, our data underlines the importance of glutathione as a player in redox homeostasis not only in the cytosol, but also in oxidative cellular compartments and it shows that its role in oxidative protein folding did already evolve in bacteria.

## Acknowledgements

We thank Franz Narberhaus for use of the fluorescence microscope for visualization of the correct localization of the roGFP probes.

## Author contributions

LRK and LIL designed the study. LRK planned and performed most of the experiments. The yeast cell assays were designed and conducted by JZ and BM. JFS performed initial experiments, established the re-oxidation assay and constructed the pPT and pPT_*roGFP2* plasmids. NL and BC purified proteins and BC assisted with DsbA *in vitro* assays. LRK and LIL wrote the manuscript. All authors consulted on the manuscript and approved the final version.

## Funding

LIL acknowledges funding from the German Research Foundation (DFG) through grant LE2905-2 and additional funding through the InnovationsFoRUM Host-Microbe-Interaction IF-018N-22-TP8.

### Data availability statement

The data supporting the findings of this study is presented within the article and its supplementary materials. Strains and plasmids constructed for this study are available upon request.

## Supplementary material

**Supplementary table 1.** . **Bacterial and yeast strains used in this study.**

| Strain | Relevant characteristics | Reference |
| --- | --- | --- |
| *E. coli* strains |  |  |
| DH5α | Cloning host | (Taylor et al., 1993) |
| BL21 (DE3) | Expression host | (Studier and Moffatt, 1986) |
| MG1655 | Expression host | (Blattner et al., 1997) |
| *E. coli* KEIO strains |  |  |
| WT (BW25113) | $\Delta(araD-araB)567$, $\Delta lacZ4787(::rrnB-3)$, $\lambda^-$, *rph*-1, $\Delta(rhaD-rhaB)568$, *hsdR*514 | (Baba et al., 2006) |
| $\Delta gshA$ (JW2663) | BW25113; $\Delta gshA769::km$ | |
| $\Delta dsbA$ (JW3832) | BW25113; $\Delta dsbA723::km$ | |
| $\Delta gshA\Delta dsbA$ | BW25113; $\Delta gshA$; $\Delta dsbA723::km$ | This study |
| $\Delta dsbB$ (JW5182-1) | BW25113; $\Delta dsbB774::km$ | (Baba et al., 2006) |
| Yeast strains |  |  |
| YPH499 | $\Delta glr1\Delta grx1\Delta grx2$ | (Zimmermann et al., 2021) |

**Supplementary table 2.**
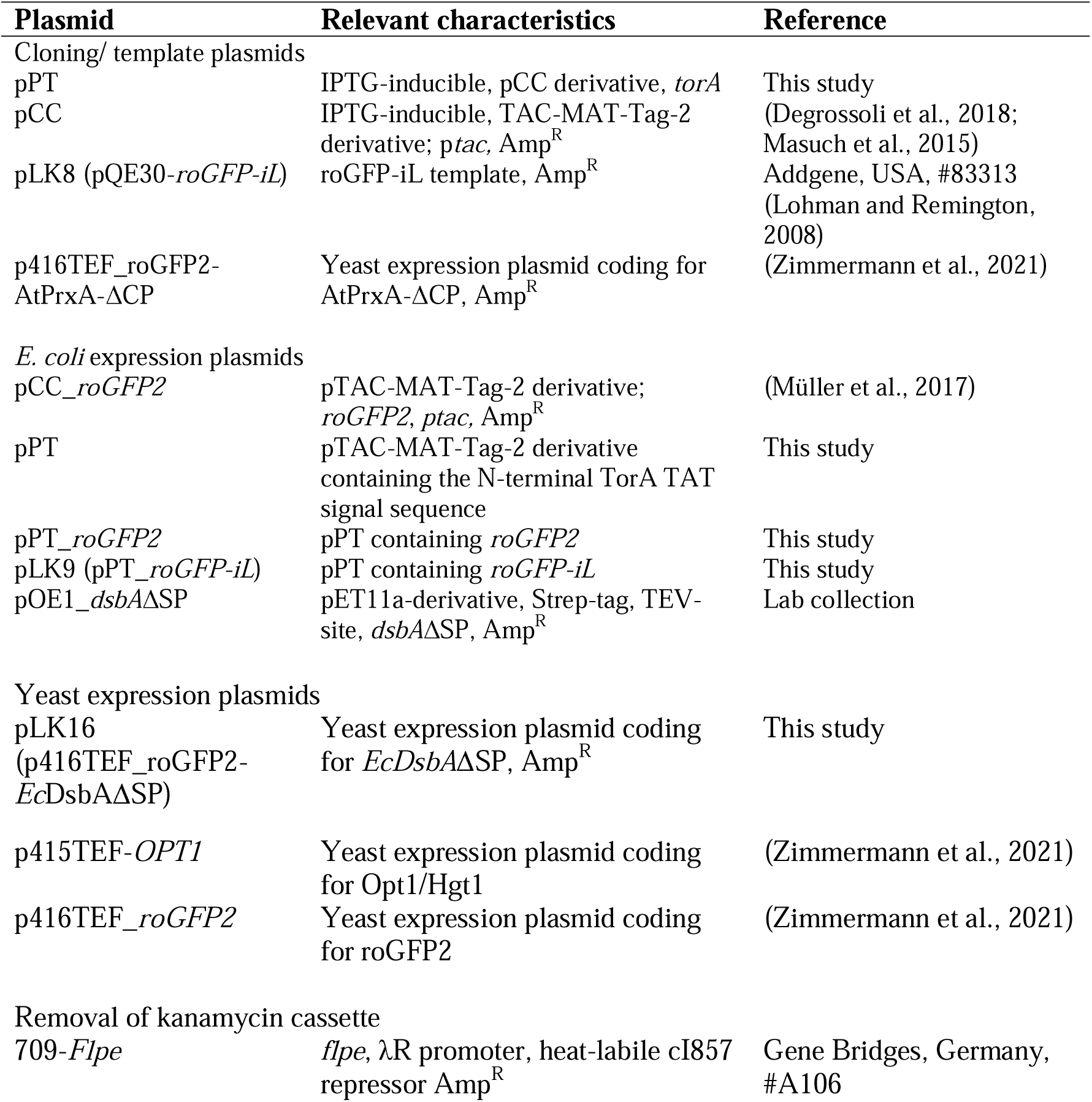
**Plasmids used in this study.**

**Supplementary table 3.**
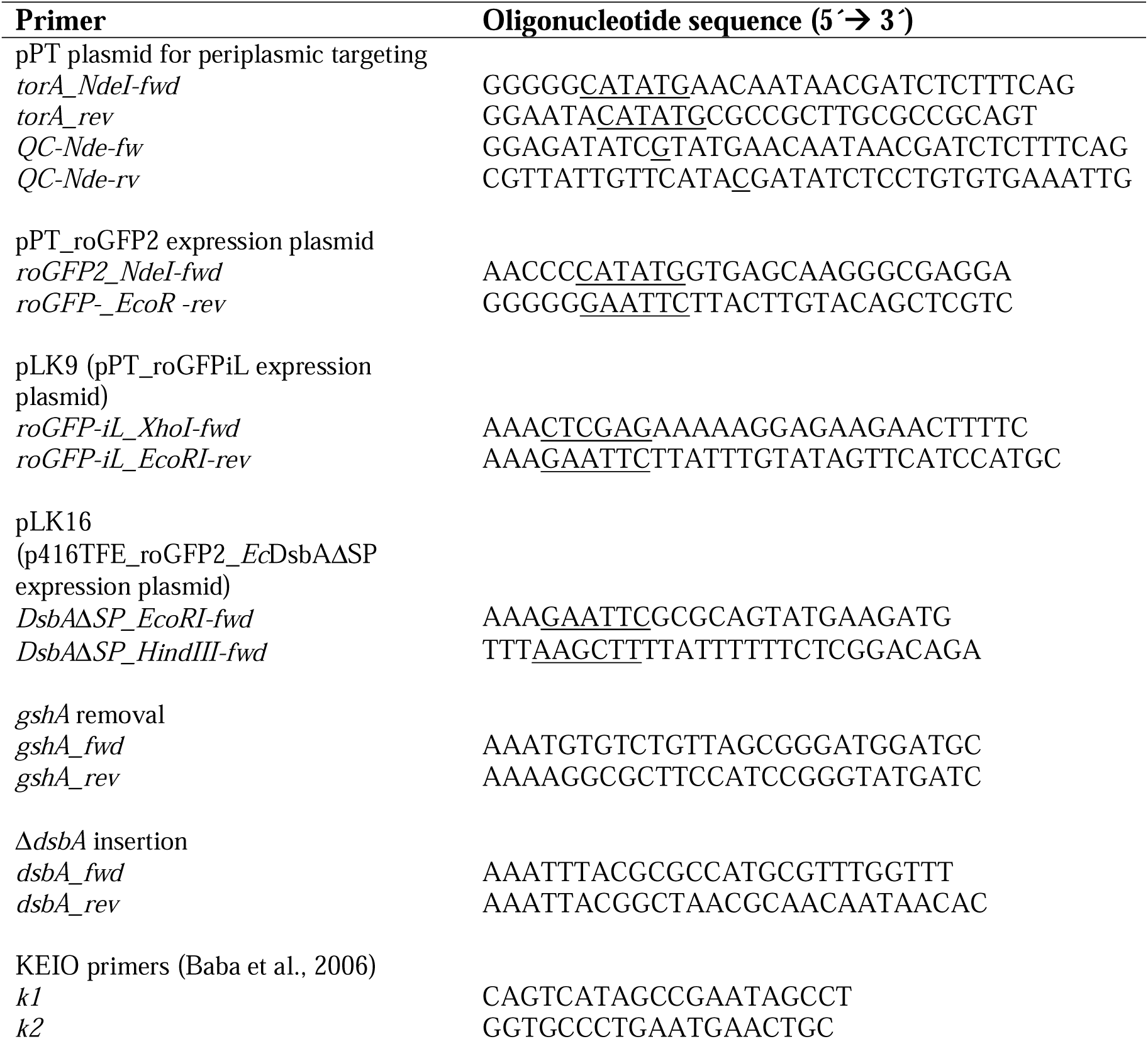
**Oligonucleotides used in this study. Restriction and mutagenesis sites are underlined.**

**Supplementary figure 2.**
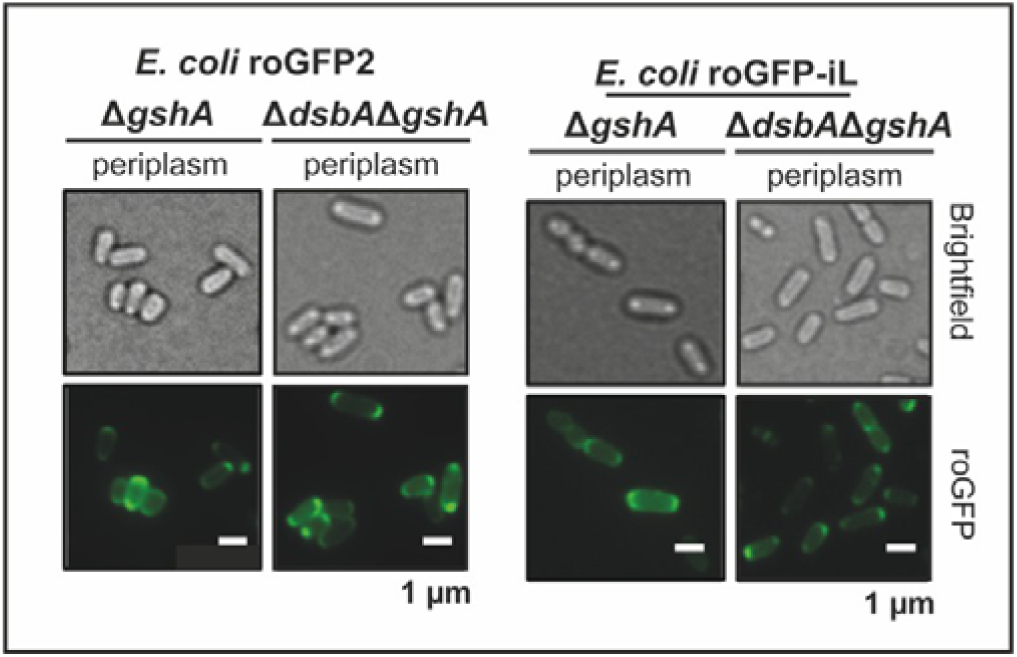
roGFP probes localize in the periplasm of cells lacking GshA or DsbA and GshA. Fluorescence microscopy of *E. coli* Δ*gshA* and Δ*dsbA*Δ*gshA* expressing roGFP2 (left) or roGFP-iL (right) confirming periplasmic localization of both roGFP probes.

**Supplementary figure 3.**
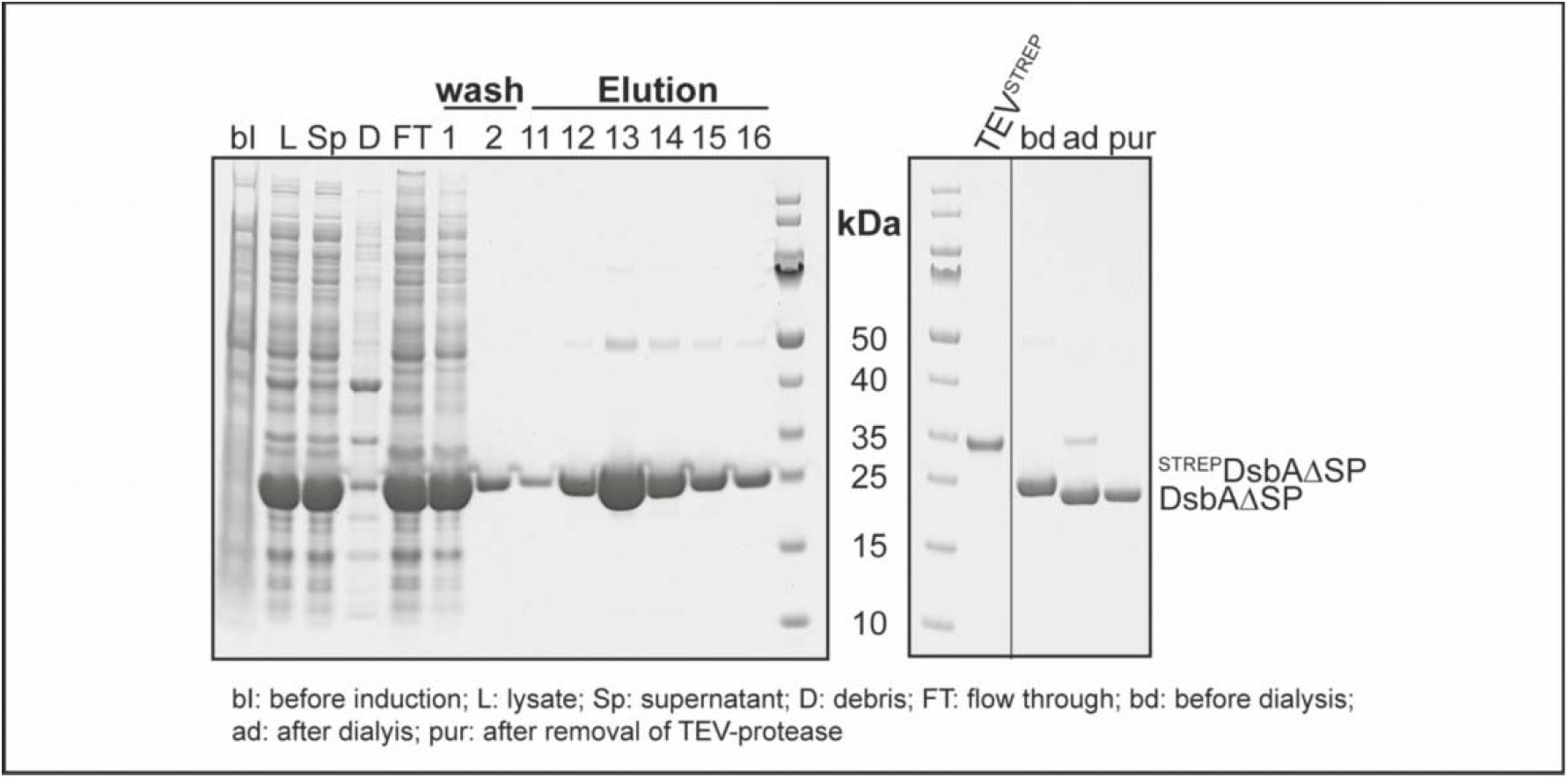
Purification of tag-free DsbAΔSP from *E. coli* BL21 cell lysate. SDS-PAGE of samples collected during the purification process. Cells harboring the plasmid for expression of ^STREP^DsbAΔSP were grown to an OD_600_ of 0.6-0.8 (bI) before expression was induced for 20 h at 20 °C. Then cells were harvested and lysed (L) and the lysate was separated by centrifugation (30 min, 4 °C, 25.000 x g) into supernatant (Sp) and debris (D). The supernatant was loaded onto a Streptavidin column (StrepTrap^TM^, GE Healthcare, Chicago, USA), washed with 5 CV buffer W (100 mM Tris-HCl, 150 mM NaCl, 1 mM EDTA, pH 8.0) and eluted with buffer E (100 mM Tris-HCl, 150 mM NaCl, 1 mM EDTA, 2.5 mM Desthiobiotin, pH 8.0) with the help of ÄKTApurifier (GE-Healthcare, Chicago, USA). Elution fractions were incubated over night at 4 °C during dialysis to buffer W with Strep-tagged TEV-protease (1:20 TEV^STREP^: ^STREP^DsbAΔSP) to get tag-free DsbA. The TEV-protease and uncleaved ^STREP^DsbAΔSP were removed using a Streptavidin column resulting in pure DsbAΔSP (pur).

**Supplementary figure 3.**
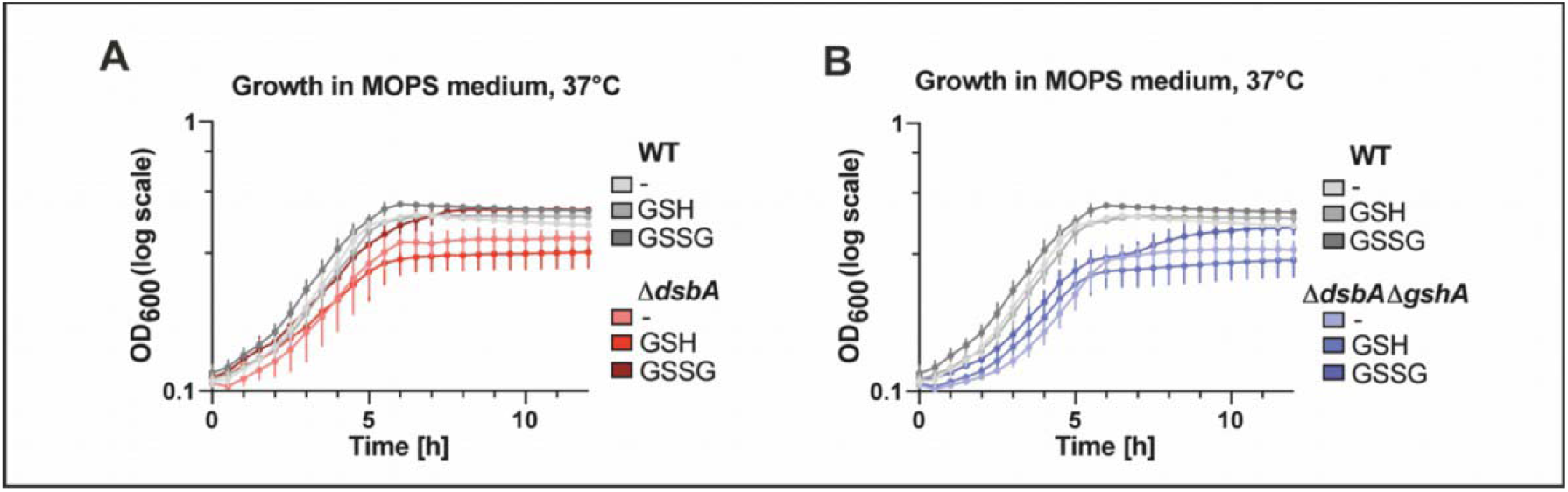
Supplementation of the growth medium with oxidized glutathione rescues growth of DsbA-deficient cells, while GSH has an inhibitory effect. *E. coli* WT (A and B), Δ*dsbA*. (**A**) and Δ*gshA*Δ*dsbA* (**B**) were cultivated at 37 °C in MOPS minimal medium without or with supplementation of 5 mM GSH or GSSG. OD_600_ was recorded over time and for better visualization y-axis is shown in log scale. Presented values are the mean of three biological replicates recorded in duplicate assays and error bars represent the standard deviation.

**Supplementary figure 4.**
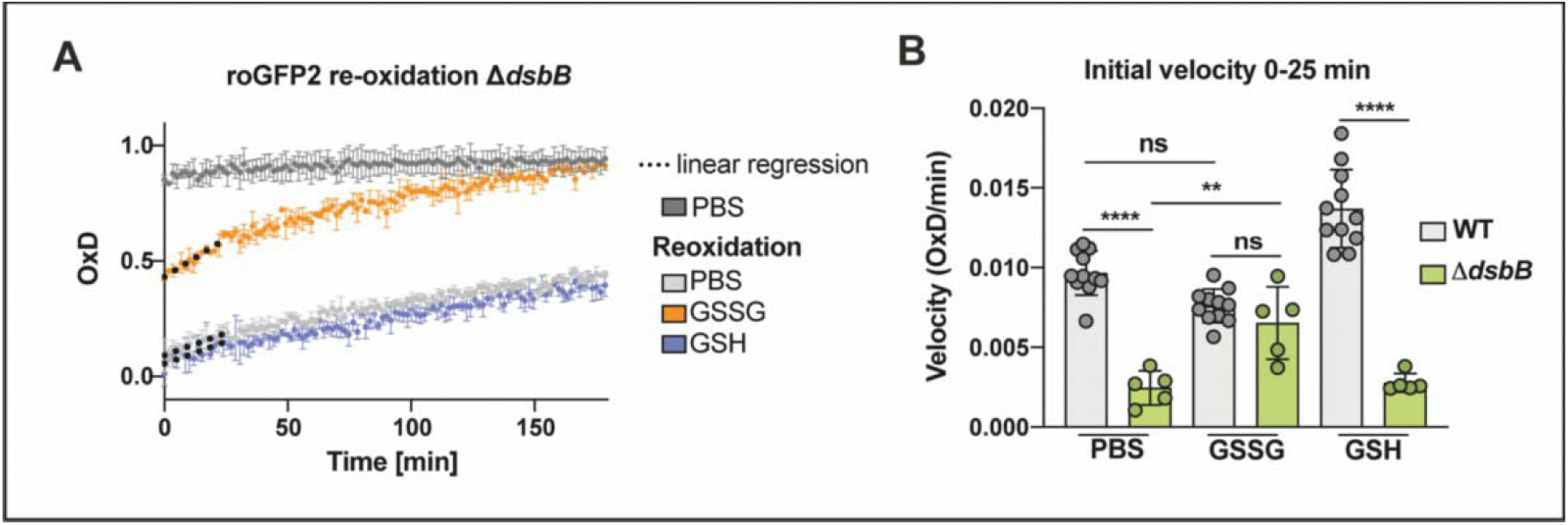
External addition of oxidized glutathione rescues the reoxidation of roGFP2 in the periplasm of cells lacking DsbB. **(A)** Reoxidation of periplasmic roGFP2 in *E. coli* Δ*dsbB*. Periplasmic reoxidation assay of roGFP2 in *E. coli* Δ*dsbB* was carried out as described in Fig. 3. One representative example out of at least five individual repeats is shown. Error bars represent standard deviation of technical triplicates. **(B)** The initial reoxidation velocity was calculated from linear regression in the first 25 min (dashed lines). Values (circles) are the mean of three technical replicates recorded in a minimum of five independent repeats. Error bars represent the standard deviation. Significance test was performed using one way ANOVA. \*\**p*<0.01, \*\*\*\**p* < 0.0001.

